# The origin of the central dogma through conflicting multilevel selection

**DOI:** 10.1101/515767

**Authors:** Nobuto Takeuchi, Kunihiko Kaneko

**Affiliations:** Research Center for Complex Systems Biology, Graduate School of Arts and Sciences, University of Tokyo, Komaba 3-8-1, Meguro-ku, Tokyo 153-8902, Japan; School of Biological Sciences, Faculty of Science, University of Auckland, Private Bag 92019, Auckland 1142, New Zealand; Department of Basic Science, Graduate School of Arts and Sciences, University of Tokyo, Komaba 3-8-1, Meguro-ku, Tokyo 153-8902, Japan

**Keywords:** reproductive division of labour, origin of genetic information, RNA world hypothesis, prebiotic evolution, Price equation

## Abstract

The central dogma of molecular biology rests on two kinds of asymmetry between genomes and enzymes: informatic asymmetry, where information flows from genomes to enzymes but not from enzymes to genomes; and catalytic asymmetry, where enzymes provide chemical catalysis but genomes do not. How did these asymmetries originate? Here we show that these asymmetries can spontaneously arise from conflict between selection at the molecular level and selection at the cellular level. We developed a model consisting of a population of protocells, each containing a population of replicating catalytic molecules. The molecules are assumed to face a trade-off between serving as catalysts and serving as templates. This trade-off causes conflicting multilevel selection: serving as catalysts is favoured by selection between protocells, whereas serving as templates is favoured by selection between molecules within protocells. This conflict induces informatic and catalytic symmetry breaking, whereby the molecules differentiate into genomes and enzymes, establishing the central dogma. We show mathematically that the symmetry breaking is caused by a positive feedback between Fisher’s reproductive values and the relative impact of selection at different levels. This feedback induces a division of labour between genomes and enzymes, provided variation at the molecular level is sufficiently large relative to variation at the cellular level, a condition that is expected to hinder the evolution of altruism. Taken together, our results suggest that the central dogma is a logical consequence of conflicting multilevel selection.

## 1 Introduction

At the heart of living systems lies a distinction between genomes and enzymes—a division of labour between the transmission of genetic information and the provision of chemical catalysis. This distinction rests on two types of asymmetry between genomes and enzymes: informatic asymmetry, where information flows from genomes to enzymes but not from enzymes to genomes; and catalytic asymmetry, where enzymes provide chemical catalysis but genomes do not. These two asymmetries constitute the essence of the central dogma in functional terms [1].

However, current hypotheses about the origin of life posit that genomes and enzymes were initially undistinguished, both embodied in a single type of molecule, RNA or its analogues [2]. While these hypotheses resolve the chicken-and-egg paradox of whether genomes or enzymes came first, they raise an obvious question: How did the genome-enzyme distinction originate?

To address this question, we explore the possibility that the genome-enzyme distinction arose during the evolutionary transition from replicating molecules to protocells [3–6]. During this transition, competition occurred both between protocells and between molecules within protocells. Consequently, selection operated at both cellular and molecular levels, and selection at one level was potentially in conflict with selection at the other [7, 8]. Previous studies have demonstrated that such conflicting multilevel selection can induce a partial and primitive distinction between genomes and enzymes in replicating molecules [9, 10]. Specifically, the molecules undergo catalytic symmetry breaking between their complementary strands, whereby one strand becomes catalytic and the other becomes non-catalytic. However, the molecules do not undergo informatic symmetry breaking—i.e., one-way flow of information from non-catalytic to catalytic molecules—because complementary replication necessitates both strands to be replicated. Therefore, the previous studies have left the most essential aspect of the central dogma unexplained.

Here we investigate whether conflicting multilevel selection can induce both informatic and catalytic symmetry breaking in replicating molecules. To this end, we extend the previous model by considering two types of replicating molecules, denoted by P and Q. Although P and Q could be interpreted as RNA and DNA, their chemical identity is unspecified for simplicity and generality. To examine the possibility of spontaneous symmetry breaking, we assume that P and Q initially do not distinguish each other. We then ask whether evolution creates a distinction between P and Q such that information flows irreversibly from one type (either P or Q) that is non-catalytic to the other that is catalytic.

## 2 Model

Our model is an agent-based model with two types of replicators, P and Q. We assume that both P and Q are initially capable of catalysing four reactions at an equal rate: the replication of P, replication of Q, transcription of P to Q, and transcription of Q to P, where complementarity is ignored (Fig. 1a; note that this figure does not depict a two-member hypercycle because in our model replicators undergo transcription [11]; see Discussion for more on comparison with hypercycles).

**Figure 1:**
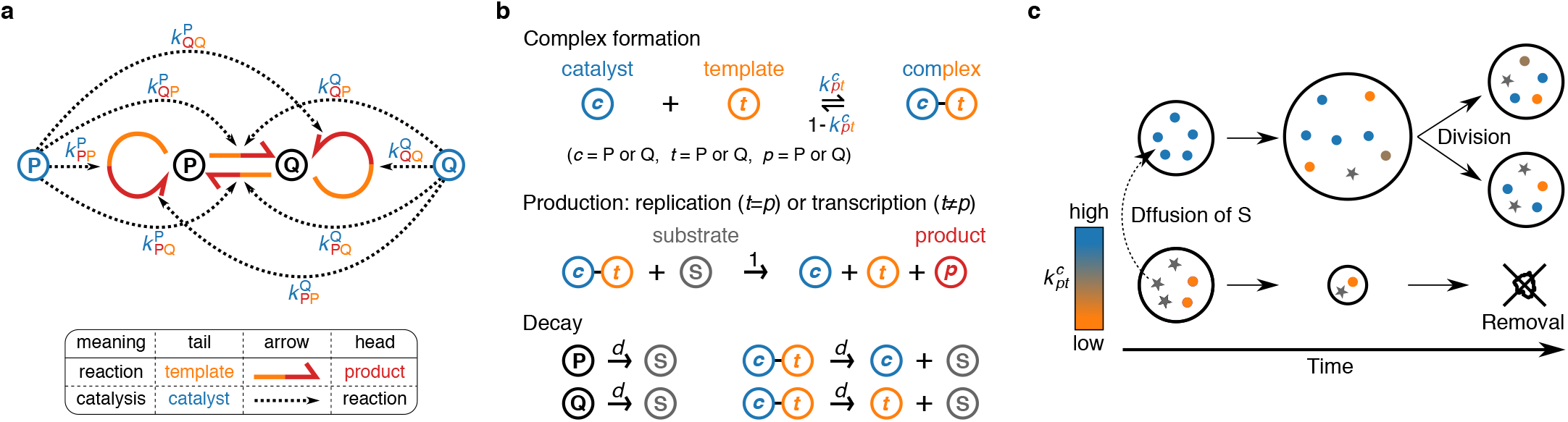
The agent-based model (see Methods for the details). **a**, Two types of replicators, P and Q, can serve as templates and catalysts for producing either type. Circular harpoons indicate replication; straight harpoons, transcription (heads indicate products; tails, templates). Dotted arrows indicate catalysis (heads indicate reaction catalysed; tails, replicators providing catalysis). **b**, Replicators undergo complex formation, replication, transcription, and decay. Rate constants of complex formation are given by the 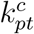 values of a replicator serving as a catalyst (whose type, P or Q, is denoted by *c*). The catalyst can form two distinct complexes with another replicator serving as a template (whose type is denoted by *t*) depending on whether it replicates (*p* = *t*) or transcribes (*p* ≠ *t*) the template. **c**, Protocells exchange substrate (represented by stars) through rapid diffusion. Protocells divide when the number of internal particles exceeds *V*. Protocells are removed when they lose all particles.

Replicators compete for a finite supply of substrate denoted by S (hereafter, P, Q, and S are collectively called particles). S is consumed through the replication and transcription of P and Q, and recycled through the decay of P and Q (Fig. 1b). Thus, the total number of particles, i.e., the sum of the total numbers of P, Q, and S is kept constant (the relative frequencies of P, Q, and S are variable).

All particles are compartmentalised into protocells, across which P and Q do not diffuse at all, but S diffuses rapidly (Fig. 1c; Methods). This difference in diffusion induces the passive transport of S from protocells in which S is converted into P and Q slowly, to protocells in which this conversion is rapid. Consequently, the latter grow at the expense of the former [12]. If the number of particles in a protocell exceeds threshold *V*, the protocell is divided with its particles randomly distributed between the two daughter cells; conversely, if this number decreases to zero, the protocell is discarded.

Crucial in our modelling is the incorporation of a trade-off between a replicator’s catalytic activities and templating opportunities. This trade-off arises from the constraint that providing catalysis and serving as a template impose structurally-incompatible requirements on replicators [13, 14]. Because replication or transcription takes a finite amount of time, serving as a catalyst comes at the cost of spending less time serving as a template, thereby inhibiting replication of itself. To incorporate this trade-off, the model assumes that replication and transcription entail complex formation between a catalyst and template (Fig. 1b) [15]. The rate constants of complex formation are given by the catalytic activities (denoted by 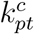) of replicators, as described below.

Each replicator is individually assigned eight catalytic values denoted by 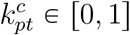, where the indices (*c*, *p*, and *t*) are P or Q (Fig. 1a). Four of these 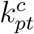 values denote the catalytic activities of the replicator itself; the other four, those of its transcripts. For example, if a replicator is of type P, its catalytic activities are given by its 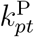 values, whereas those of its transcripts, which are of type Q, are given by its 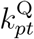 values. The indices *p* and *t* denote the specific type of reaction catalysed, as depicted in Fig. 1a. When a new replicator is produced, its 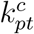 values are inherited from its template with potential mutation of probability *m* (Methods).

The 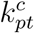 values of a replicator determine the rates at which this replicator forms a complex with another replicator and catalyses replication or transcription of the latter (Fig. 1b; Methods). The greater the catalytic activities 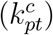 of a replicator, the greater the chance that the replicator is sequestered in a complex as a catalyst and thus unable to serve as a template—hence a trade-off. Note that the trade-off is relative: if all replicators in a protocell have identical 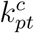 values, their multiplication rate increases monotonically with their 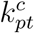 values, assuming all else is held constant.

The above trade-off creates a dilemma: providing catalysis brings benefit at the cellular level because it accelerates a protocell’s uptake of substrate; however, providing catalysis brings cost at the molecular level because it decreases the relative opportunity of a replicator to be replicated within a protocell [9]. Therefore, selection between protocells tends to maximise the 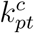 values of replicators (i.e., cellular-level selection), whereas selection within protocells tends to minimise the 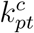 values of replicators (i.e., molecular-level selection).

## 3 Results

### 3.1 Computational analysis

Using the agent-based model described above, we examined how 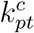 values evolve as a result of conflicting multilevel selection. To this end, we set the initial 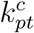 values of all replicators to 1, so that P and Q are initially identical in their catalytic activities (the initial frequencies of P or Q are also set to be equal). We then simulated the model for various values of *V* (the threshold at which protocells divide) and *m* (mutation rate).

Our main result is that for sufficiently large values of *V* and *m*, replicators undergo spontaneous symmetry breaking in three aspects (Figs. 2a-d and S1). First, one type of replicator (either P or Q) evolves high catalytic activity, whereas the other completely loses it (i.e., 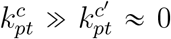 for *c* ≠ *c*′): catalytic symmetry breaking (Fig. 2bc). Second, templates are transcribed into catalysts, but catalysts are not reverse-transcribed into templates (i.e., 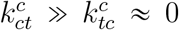): informatic symmetry breaking (Fig. 2bc). Finally, the copy number of templates becomes smaller than that of catalysts: numerical symmetry breaking: (Fig. 2d). This three-fold symmetry breaking is robust to various changes in model details (see SI Text 1.1 and 1.2; Figs. S2, S3, and S4).

**Figure 2:**
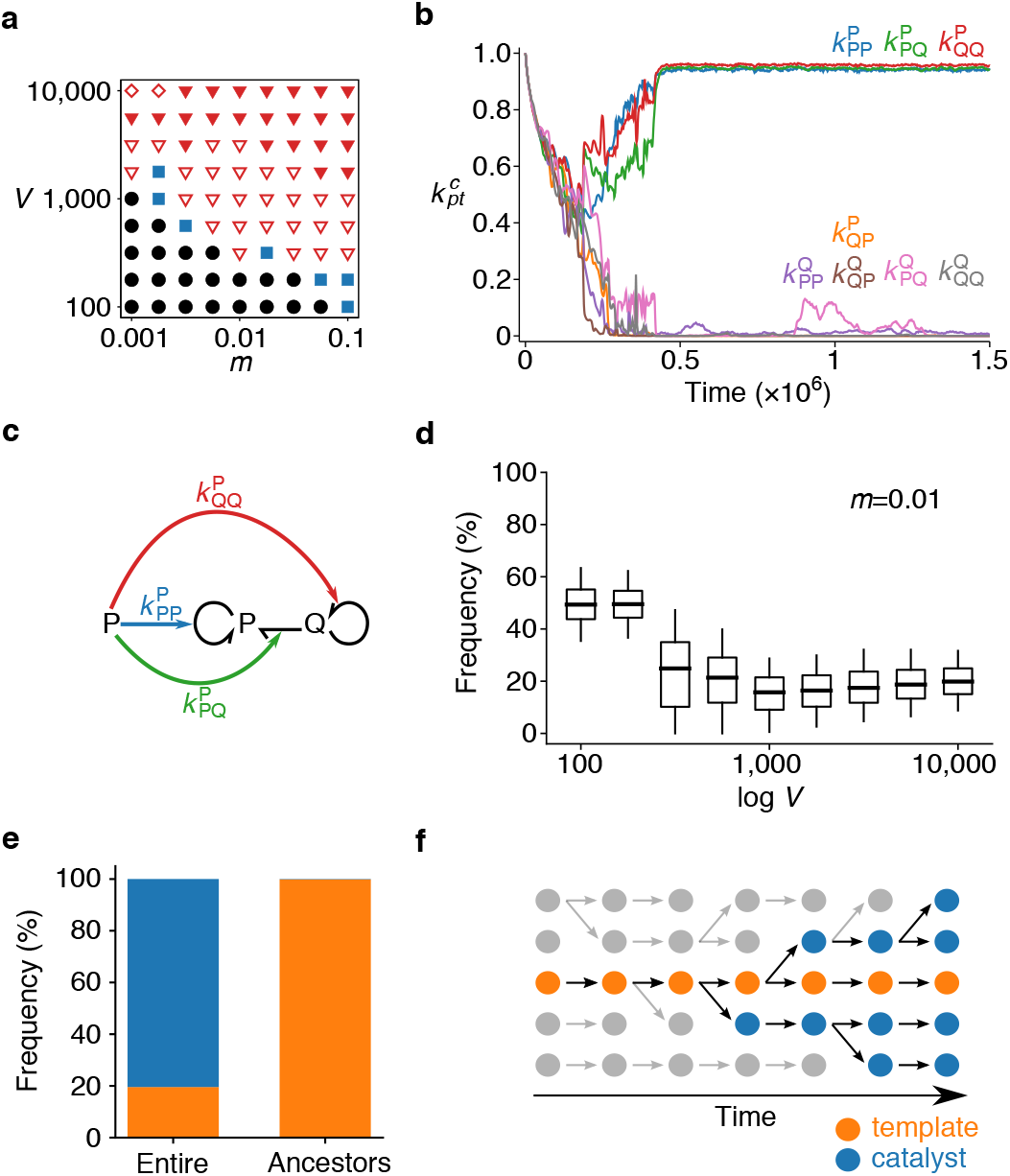
The evolution of the central dogma. **a**, Phase diagram: circles indicate no symmetry breaking (Fig. S1ab); squares, uncategorised (Fig. S1cd); open triangles, incomplete symmetry breaking (Fig. S1e-h); filled triangles, three-fold symmetry breaking as depicted in b, c, and d; diamonds, catalytic and informatic symmetry breaking without numerical symmetry breaking (Fig. S5a). The initial condition was 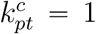 for all replicators. **b**, Dynamics of 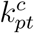 averaged over all replicators. *V* = 10000 and *m* = 0.01. **c**, Catalytic activities evolved in b. **d**, Per-cell frequency of minority replicator types (P or Q) at equilibrium as a function of *V*: boxes, quartiles; whiskers, 5th and 95th percentiles. Only protocells containing at least *V*/2 particles were considered. **e**, Frequencies of templates (orange) and catalysts (blue) in the entire population or in the common ancestors. *V* = 3162 and *m* = 0.01. **f**, Illustration of e. Circles represent replicators; arrows, genealogy. Extinct lineages are grey. Common ancestors are always templates, whereas the majority of replicators are catalysts.

A significant consequence of the catalytic and informatic symmetry breaking is the resolution of the dilemma between providing catalysis and getting replicated. Once symmetry is broken, tracking lineages reveals that the common ancestors of all replicators are almost always templates (Fig. 2ef; Methods). That is, information is transmitted almost exclusively through templates, whereas information in catalysts is eventually lost (i.e., catalysts have zero reproductive value). Consequently, evolution operates almost exclusively through competition between templates, rather than between catalysts. How the catalytic activity of catalysts evolves, therefore, depends solely on the cost and benefit to templates. On one hand, this catalytic activity brings benefit to templates for competition across protocells. On the other hand, this activity brings no cost to templates for competition within a protocell (neither does it bring benefit because catalysis is equally shared among templates). Therefore, the catalytic activity of catalysts is maximised by cellular-level selection operating on templates, but not minimised by molecular-level selection operating on templates, hence the resolution of the dilemma between catalysing and templating. Because of this resolution, symmetry breaking leads to the maintenance of high catalytic activities (Figs. S6 and S7).

### 3.2 Mathematical analysis

To understand the mechanism of the catalytic and informatic symmetry breaking, we simplified the agent-based model into mathematical equations. These equations allow us to consider all the costs and benefits involved in the provision of catalysis by *c* ∈ {P, Q}: molecular-level cost to *c* (denoted by 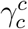) and cellular-level benefit to *t* ∈ {P, Q} (denoted by 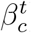). The equations calculate the joint effects of all these costs and benefits on the evolution of the average catalytic activities of *c* (denoted by 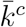). The equations are derived with the help of Price’s theorem [7, 8, 16] and displayed below (see Methods and SI Text 1.3 for the derivation):

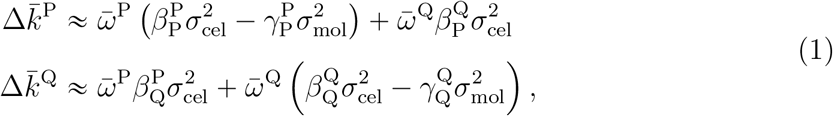

where Δ denotes evolutionary change per generation, 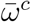 is the average normalised reproductive value of *c*, 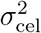 is the variance of catalytic activities among protocells (cellular-level variance), and 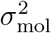 is the variance of catalytic activities within a protocell (molecular-level variance).

The first and second terms on the right-hand side of equations (1) represent evolution arising through the replication of P and Q, respectively, weighted by the reproductive values, 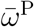 and 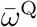. The terms multiplied by 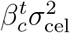 represent evolution driven by cellular-level selection; those by 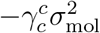, evolution driven by molecular-level selection.

The derivation of equations (1) involves various simplifications that are not made in the agent-based model, among which the three most important are noted below (see Methods and SI Text 1.3 for details). First, equations (1) simplify evolutionary dynamics by restricting the number of evolvable parameters to a minimum required for catalytic and informatic symmetry breaking. More specifically, equations (1) assume that 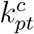 is independent of *p* and *t* (denoted by *k*^*c*^), i.e., catalysts do not distinguish the replicator types of templates and products. Despite this simplification, catalytic symmetry breaking can still occur (e.g., *k*^P^ > *k*^Q^), as can informatic symmetry breaking: the trade-off between catalysing and templating causes information to flow preferentially from less catalytic to more catalytic replicator types. However, numerical symmetry breaking is excluded as it requires 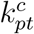 to depend on *p*; consequently, the frequencies of P or Q are fixed and even in equations (1) (this is not the case in the agent-based model described in the previous section). Therefore, while equations (1) are useful for identifying the mechanism of catalytic and informatic symmetry breaking, they are not useful for identifying the mechanism of numerical symmetry breaking. In a supplementary material, we use different equations to identify the mechanism of numerical symmetry breaking (see SI Text 1.4 and Fig. S5).

The second simplification involved in equations (1) is that variances 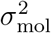 and 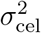 are treated as parameters although they are actually dynamic variables dependent on *m* and *V* in the agent-based model (in supplementary material, we examine this assumption; see SI Text 1.5 and Fig. S8). In addition, these variances are assumed to be identical between 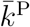 and 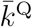 because no difference is a priori assumed between P and Q.

The third simplification involved in equations (1) is that the terms of order greater than 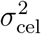 and 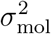 are ignored under the assumption of weak selection [16].

Using equations (1), we can now elucidate the mechanism of the symmetry breaking. Consider a symmetric situation where P and Q are equally catalytic: 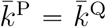. Since P and Q are identical, the catalytic activities of P and Q evolve identically: 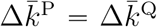. Next, suppose that P becomes slightly more catalytic than Q for whatever reason, e.g., by genetic drift: 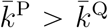 (catalytic asymmetry). The trade-off between catalysing and templating then causes P to be replicated less frequently than Q, so that 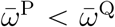 (informatic asymmetry). Consequently, the second terms of equations (1) increase relative to the first terms. That is, for catalysis provided by P (i.e., 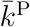), the impact of cellular-level selection through Q (i.e., 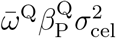) increases relative to those of molecular-level and cellular-level selection through P (i.e., 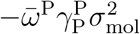 and 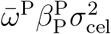, respectively), resulting in the relative strengthening of cellular-level selection. By contrast, for catalysis provided by Q (i.e., 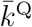), the impacts of molecular-level and cellular-level selection Q (i.e., 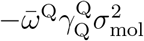 and 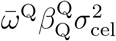, respectively) increase relative to cellular-level selection through through P (i.e., 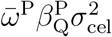), resulting in the relative strengthening of molecular-level selection. Consequently, a small difference between 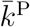 and 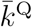 leads to 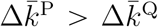, the amplification of the initial difference—hence, symmetry breaking. The above mechanism can be summarised as a positive feedback between reproductive values and the relative impact of selection at different levels.

We next asked whether, and under what conditions, the above feedback leads to symmetry breaking such that either P or Q completely loses catalytic activity. To address this question, we performed a phase-plane analysis of equations (1) as described in Fig. 3 (see Methods and SI Text 1.6 for details). Figure 3 shows that 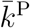 and 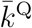 diverge from symmetric states (i.e., 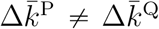), confirming the positive feedback described above. However, symmetry breaking occurs only if molecular-level variance 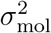 is sufficiently large relative to cellular-level variance 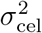 [i.e., if genetic relatedness between replicators, 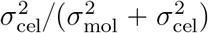, is sufficiently low; see Methods]. Large 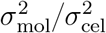 is required because if 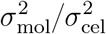 is too small, cellular-level selection completely dominates over molecular-level selection, maximising both 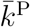 and 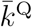 (Fig. 3a). The requirement of large 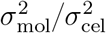 is consistent with the fact that the agent-based model displays symmetry breaking for sufficiently large *V*: the law of large numbers implies that 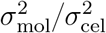 increases with *V* [9, 17]. This consistency with the agent-model suggests that equations (1) correctly describe the mechanism of symmetry breaking in the agent-based model (see SI Text 1.5 and Fig. S8 for an additional consistency check in terms of both *m* and *V*).

**Figure 3:**
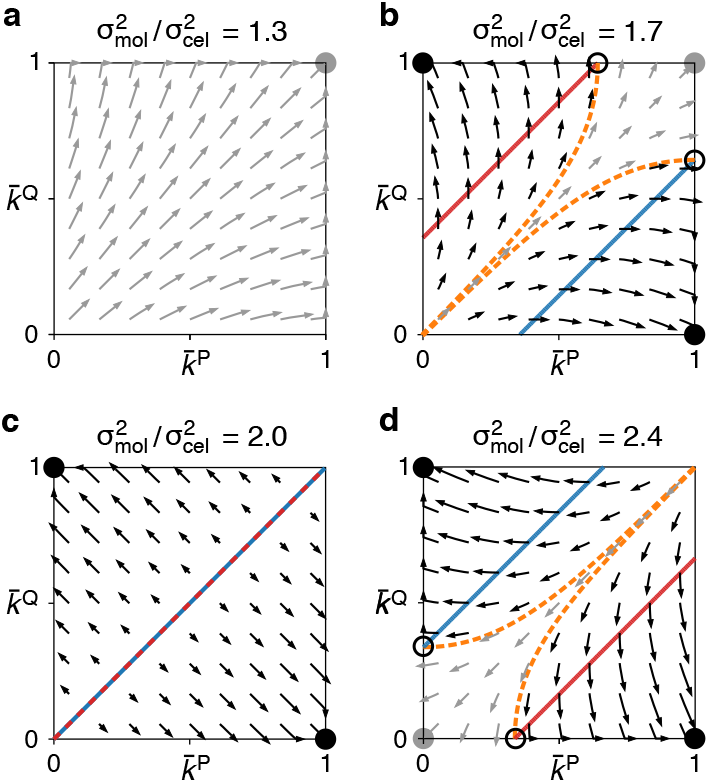
Phase-plane analysis. For this analysis, equations (1) were adapted as follows: 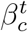 and 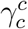 were set to 1; 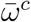 was calculated as 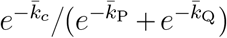; Δ was replaced with time derivative 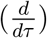; and 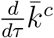 was set to 0 if 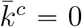 or 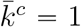 to ensure that 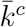 is bounded within [0, 1] as in the agent-based model. Solid lines indicate nullclines: 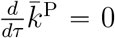 (red) and 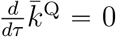 (blue). The nullclines at 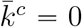 and 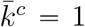 are not depicted for visibility. Filled circles indicate symmetric (grey) and asymmetric (black) stable equilibria; open circles, unstable equilibria; arrows, short-duration flows (Δ*τ* = 0.15) leading to symmetric (grey) or asymmetric (black) equilibria. Dashed lines (orange) demarcate basins of attraction. 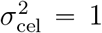. **a**, Molecular-level variance is so small that cellular-level selection completely dominates; consequently, 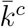 is always maximised. **b**, Molecular-level variance is large enough to create asymmetric equilibria; however, cellular-level variance is still large enough to make 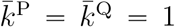 stable. **c**, A tipping point; the nullclines overlap. **d**, Molecular-level variance is so large that 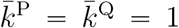 is unstable; the asymmetric equilibria can be reached if 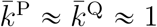.

## 4 Discussion

Our results show that conflicting multilevel selection can induce informatic and catalytic symmetry breaking in replicating molecules. The symmetry breaking is induced because molecular-level selection minimises the catalytic activity of one type of molecule (either P or Q), whereas cellular-level selection maximises that of the other. The significance of the symmetry breaking is that it results in one-way flow of information from non-catalytic to catalytic molecules—the central dogma. The symmetry breaking thereby establishes a division of labour between the transmission of genetic information and the provision of chemical catalysis. This division of labour resolves a dilemma between templating and catalysing, the very source of conflict between levels of selection. Below, we discuss our results in relation to four subjects, namely, chemistry, hypercycle theory, kin selection theory, and reproductive division of labour.

Our theory does not specify the chemical details of replicating molecules, and this abstraction carries two implications. First, our theory suggests that the central dogma, if formulated in functional terms, is a general feature of living systems that is independent of protein chemistry. When the central dogma was originally proposed, it was formulated in chemical terms as the irreversible flow of information from nucleic acids to proteins [1]. Accordingly, the chemical properties of proteins have been considered integral to the central dogma [18]. By contrast, the present study formulates the central dogma in functional terms, as the irreversible flow of information from non-catalytic to catalytic molecules. Our theory shows that the central dogma, formulated as such, is a logical consequence of conflicting multilevel selection. Therefore, the central dogma might be a general feature of life that is independent of the chemical specifics of material in which life is embodied.

The second implication of the chemical abstraction is that our theory could be tested by experiments with existing materials. Our theory assumes that a replicator faces a trade-off between providing ‘catalysis’ and getting replicated. However, it does not restrict catalysis to being replicase activity: although our agent-based model assumes that catalysts are replicases, our mathematical analysis does not. Therefore, existing RNA and DNA molecules could be used to test our theory [19]. For example, one could compare two systems, one where RNA serves as both templates and catalysts, and one where RNA serves as catalysts and DNA serves as templates. According to our theory, the latter is expected to maintain higher catalytic activity through evolution, provided the mutation rate and the number of molecules per cell are sufficiently large (see also [20]). In addition, using RNA and DNA is potentially relevant to the historical origin of the central dogma, given the possibility that DNA might have emerged before the advent of protein translation [21–24].

While our theory is similar to hypercycle theory in that both are concerned with the evolution of complexity in replicator systems, our theory proposes a distinct mechanism for evolving such complexity. Whereas hypercycle theory proposes symbiosis between multiple lineages of replicators [11], our theory proposes symmetry breaking (i.e., differentiation) in a single lineage of replicators—a fundamental distinction that is drawn between ‘egalitarian’ and ‘fraternal’ major evolutionary transitions as defined by Queller [25] (egalitarianism implies equality, which is involved in the evolution of complexity through symbiosis, whereas fraternalism implies kinship, which is involved in the evolution of complexity through differentiation; these terms are taken from the French Revolutionary slogan, *‘Liberté, Egalité, Fraternité’*).

Moreover, our theory differs from hypercycle theory in terms of the roles played by non-catalytic templates. In hypercycle theory, the evolution of non-catalytic templates jeopardises hypercycles because such templates (called parasites) can replicate faster than catalytic templates constituting the hypercycles [15, 26]. In our theory, the evolution of non-catalytic templates is one of the essential factors leading to the division of labour between genomes and enzymes.

While our theory differs from hypercycle theory in the above aspects, it does not contradict the latter. In fact, there is a potential synergy between the evolution of complexity through symmetry breaking and that through symbiosis. Our theory posits that a distinction between genomes and enzymes resolves the dilemma between templating and catalysing, thereby increasing the evolutionary stability of catalytic activities in replicators. Likewise, this distinction might also contribute to the evolutionary stability of symbiosis between replicators, hence the potential synergy (however, we should add that the specific mechanism of symbiosis proposed by hypercycle theory is not unique [27–33]).

While our theory is consistent with kin selection theory, it makes a novel prediction for evolution under a condition of low genetic relatedness. Kin selection theory posits that altruism can evolve if genetic relatedness is sufficiently high [34]. Consistent with this, our theory posits that for sufficiently high genetic relatedness (i.e., for sufficiently high 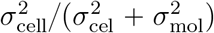, or sufficiently small *m* and *V*), cellular-level selection maximises the provision of catalysis by all molecules, establishing full altruism (providing catalysis can be viewed as altruism [35]: providing catalysis brings no direct benefit to a catalyst because a catalyst cannot catalyse the replication of itself in our model). However, the two theories diverge for sufficiently low genetic relatedness. In this case, kin selection theory predicts that evolution cannot lead to altruism; by contrast, our theory predicts that evolution can lead to a division of labour between the transmission of genetic information and the provision of chemical catalysis. Whether this reproductive division of labour should be called altruism is up for debate.

In relation to reproductive division of labour, our theory suggests a novel mechanism for its evolution in terms of a distinction between genomes and enzymes. In previous theories, reproductive division of labour has been regarded as an adaptation caused by natural selection [4–6]. For example, Michod has shown that reproductive division of labour can evolve because it maximises the group-level fitness of replicating entities if a trade-off curve between the replicating capacity and other functional capacities of the entities is convex [36] (see [37] for a historical reference). In our theory, division of labour between genomes and enzymes evolves, not because it maximises the fitness of a protocell (i.e., group), but because it is a stable equilibrium between evolution driven by molecular-level selection and evolution driven by cellular-level selection, an emergent outcome of conflicting multilevel selection (note that the fitness of a protocell is maximal if all replicators in the protocell are maximally catalytic and hence display no division of labour, a state that evolves for sufficiently small *V* and *m*). Parallel results have been obtained from previous studies, where conflicting multilevel selection is shown to evolve various states that are not directly selected for at any single level [9, 20, 38]. Taken together, these results suggest the possibility that biological complexity evolves as emergent outcomes of conflicting multilevel selection.

Finally, we note that the division of labour between the transmission of genetic information and other functions is a recurrent pattern throughout biological hierarchy. For example, multicellular organisms display differentiation between germline and soma, as do eusocial animal colonies between queens and workers (Table 1) [3–6]. Given that all these systems potentially involve conflicting multilevel selection and tend to display reproductive division of labour as their sizes increase [6], our theory might provide a basis on which to pursue a universal principle of life that transcends the levels of biological hierarchy.

**Table 1:** Division of labour between information transmission and other functions transcends the levels of biological hierarchy.

| hierarchy |  | differentiation |  |
| --- | --- | --- | --- |
| whole | parts | information | other |
| cell | molecules | genome | enzyme |
| symbiont population* | prokaryotic cells | transmitted | non-transmitted |
| ciliate | organelles | micronucleus | macronucleus |
| multicellular organism | eukaryotic cells | germ | soma |
| eusocial colony | animals | queen | worker |

\*Bacterial endosymbionts of ungulate lice (*Haematopinus*) and planthoppers (*Fulgoroidea*) [39].

## 5 Methods

### 5.1 Details of the model

The model treats each molecule as a distinct individual with uniquely-assigned 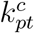 values. One time step of the model consists of three sub-steps: reaction, diffusion, and cell division.

In the reaction step, the reactions depicted in Fig. 1b are simulated with the algorithm described previously [9]. The rate constants of complex formation are given by the 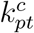 values of a replicator serving as a catalyst. For example, if two replicators, denoted by *X* and *Y*, serve as a catalyst and template, respectively, the rate constant of complex formation is the 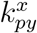 value of *X*, where *x*, *y*, and *p* are the replicator types (i.e., P or Q) of *X*, *Y*, and product, respectively. If *X* and *Y* switch the roles (i.e., *X* serves as a template, and *Y* serves as a catalyst), the rate constant of complex formation is the 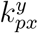 value of *Y*. Complexes are distinguished not only by the roles of *X* and *Y*, but also by the replicator type of product *p*. Therefore, *X* and *Y* can form four distinct complexes depending on which replicator serves as a catalyst and which type of replicator is being produced.

The above rule about complex formation implies that whether a template is replicated (*p* = *t*) or transcribed (*p* ≠ *t*) depends entirely on the 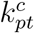 values of a catalyst. In other words, a template cannot control how its information is used by a catalyst. This rule excludes the possibility that a template maximises its fitness by biasing catalysts towards replication rather than transcription. Excluding this possibility is legitimate if the backbone of a template does not directly determine the backbone of a product as in nucleic acid polymerisation.

In addition, the above rule about complex formation implies that replicators multiply fastest if their 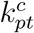 values are maximised for all combinations of *c*, *p*, and *t* (this is because *X* and *Y* form a complex at a rate proportional to 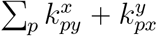 if all possible complexes are considered). Consequently, cellular-level selection tends to maximize 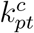 values for all combinations of *c*, *p*, and *t* (because cellular-level selection tends to maximise the multiplication rate of replicators within protocells). If 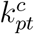 values are maximised for all combinations of *c*, *p*, and *t*, P and Q coexist. Therefore, coexistence between P and Q is favoured by cellular-level selection, a situation that might not always be the case in reality. We ascertained that the above specific rule about complex formation does not critically affect results by examining an alternative model in which cellular-level selection does not necessarily favour coexistence between P and Q (see SI Text 1.1).

In the diffusion step, all substrate molecules are randomly re-distributed among protocells with probabilities proportional to the number of replicators in protocells. In other words, the model assumes that substrate diffuses extremely rapidly.

In the cell-division step, every protocell containing more than *V* particles (i.e. P, Q, and S together) is divided as described in Model.

The mutation of 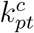 is modelled as unbiased random walks. With a probability *m* per replication or transcription, each 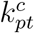 value of a replicator is mutated by adding a number randomly drawn from a uniform distribution on the interval (−*δ*_mut_, *δ*_mut_) (*δ*_mut_ = 0.05 unless otherwise stated). The values of 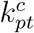 are bounded above by *k*_max_ with a reflecting boundary (*k*_max_ = 1 unless otherwise stated), but are not bounded below to remove the boundary effect at 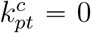. However, if 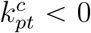, the respective rate constant of complex formation is regarded as zero.

We ascertained that the above specific model of mutation does not critically affect results by testing two alternative models of mutation. One model is nearly the same as the above, except that the boundary condition at 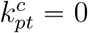 was set to reflecting. The other model implements mutation as unbiased random walks on a logarithmic scale. The details are described in SI Text 1.2.

Each simulation was run for at least 5 × 10^7^ time steps (denoted by *t*_min_) unless otherwise stated, where the unit of time is defined as that in which one replicator decays with probability *d* (thus, the average lifetime of replicators is 1/*d* time steps). The value of *d* was set to 0.02. The total number of particles in the model *N*_tot_ was set to 50*V* so that the number of protocells was approximately 100 irrespective of the value of *V*. At the beginning of each simulation, 50 protocells of equal size were generated. The initial values of 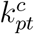 were set to *k*_max_ for every replicator unless otherwise stated. The initial frequencies of P and Q were equal, and that of S was zero.

### 5.2 Ancestor tracking

Common ancestors of replicators were obtained in two steps. First, ancestor tracking was done at the cellular level to obtain the common ancestors of all surviving protocells. Second, ancestor tracking was done at the molecular level for the replicators contained by the common ancestors of protocells obtained in the first step. The results shown in Fig. 2e were obtained from the data between 2.1 × 10^7^ and 2.17 × 10^7^ time steps, so that the ancestor distribution was from after the completion of symmetry breaking.

### 5.3 Outline of the derivation of equations (1)

To derive equations (1), we simplified the agent-based model in two ways. First, we assumed that 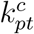 is independent of *p* and *t*. Under this assumption, a catalyst does not distinguish the replicator types of templates (i.e., 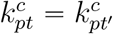 for *t* ≠ *t*′) and products (i.e., 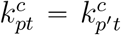 for *p* ≠ *p*′). This assumption excludes the possibility of numerical symmetry breaking, but still allows catalytic and informatic symmetry breaking as described in Results.

Second, we abstracted away chemical reactions by defining 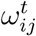 as the probability that replicator *j* of type *t* in protocell *i* is replicated or transcribed per unit time. Let 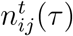 be the population size of this replicator at time *τ*. Then, 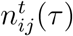 is expected to satisfy

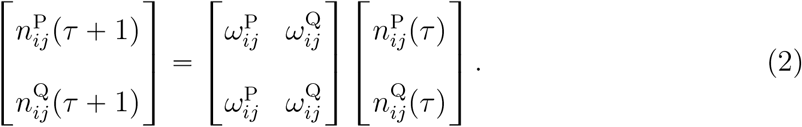

The fitness of the replicator can be defined as the dominant eigenvalue *λ*_*ij*_ of the 2 × 2 matrix on the right-hand side of equation (2): 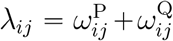. Fisher’s reproductive values of P and Q are given by the corresponding left eigenvector 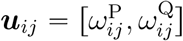.

The evolutionary dynamics of the average catalytic activity of replicators can be described with Price’s equation [7, 8]. Let 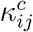 be the catalytic activity of replicator *j* of type *c* in protocell *i* (we use *κ* instead of *k* to distinguish 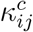 from 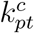). Price’s equation states that

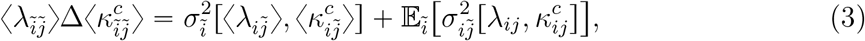

where 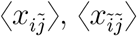, and 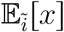 are *x* averaged over the indices marked with tildes, 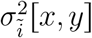 is the covariance between *x* and *y* over protocells, and 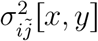 is the covariance between *x* and *y* over the replicators in protocell *i*. One replicator is always counted as one sample in calculating all moments.

To approximate equation (3), we assumed that covariances between 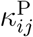 and 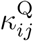 and between 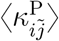 and 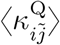 are negligible because the mutation of 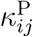 and that of 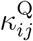 are uncorrelated in the agent-based model (see SI Text 1.6 for an alternative justification of this assumption). Under this assumption, equation (3) is approximated by equations (1) up to the second central moments of 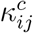 and 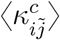, with the following notation (see SI Text 1.3 for the derivation):

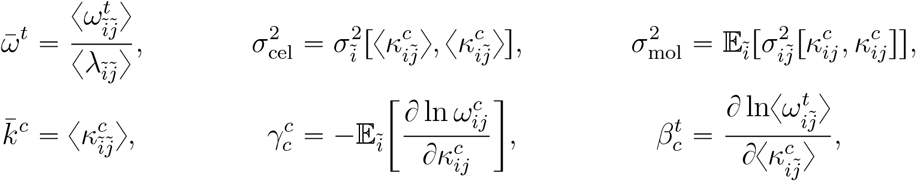

where 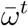 is the normalised average reproductive value of type-*t* replicators, 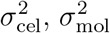, and 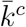 are the simplification of the notation, 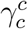 is an average decrease in the replication rate of a type-*c* replicator due to an increase in its own catalytic activity, and 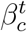 is an increase in the average replication rate of type-*t* replicators in a protocell due to an increase in the average catalytic activity of type-*c* replicators in that protocell. We assumed that 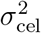 and 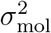 do not depend on *c* because no difference is a priori assumed between P and Q.

The values of 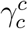 and 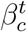 can be interpreted as the cost and benefit of providing catalysis. Let us assume that *V* is so large that 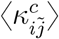 and 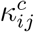 can be regarded as mathematically independent of each other if *i* and *j* are fixed (if *i* and *j* are varied, 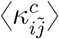 and 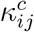 may be statistically correlated). Under this assumption, increasing 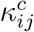 does not increase 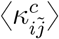, so that 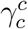 reflects only the cost of providing catalysis at the molecular level. Likewise, increasing 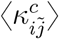 does not increase 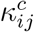, so that 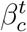 reflects only the benefit of receiving catalysis at the cellular level. Moreover, the independence of 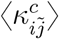 from 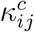 implies that 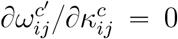 for *c* ≠ *c*′, which permits the following interpretation: if a replicator of type *c* provides more catalysis, its transcripts, which is of type *c*′, pay no extra cost (i.e., 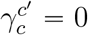).

### 5.4 Outline of the phase-plane analysis

To perform the phase-plane analysis depicted in Fig. 3, we defined 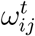 as a specific function of 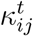 (see above for the meaning of 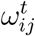 and 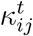):

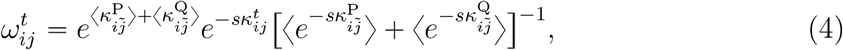

where the first factor 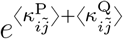 represents the cellular-level benefit of catalysis provided by the replicators in protocell *i*, the second factor 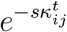 represents the molecular-level cost of catalysis provided by the focal replicator, the last factor normalises the cost, and *s* is the cost-benefit ratio. The above definition of 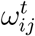 was chosen to satisfy the requirement that a replicator faces the trade-off between providing catalysis and serving as a template, i.e., 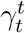 and 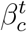 are positive. Apart from this requirement, the definition was arbitrarily chosen for simplicity.

Under the definition in equation (4), we again approximated equation (3) up to the second central moments of 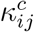 and 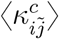, obtaining the following (see SI Text 1.6 for the derivation):

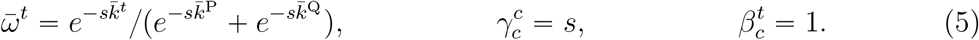

Equations (1) and (5) can be expressed in a compact form as

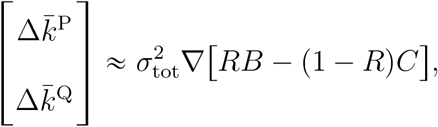

where 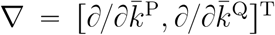 (^T^ denotes transpose), 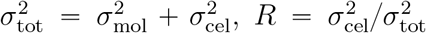, 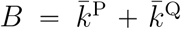, and 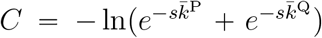. *R* can be interpreted as the regression coefficient of 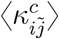 on 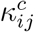 [40] and, therefore, the coefficient of genetic relatedness [41]. The potential *RB* − (1 − *R*)*C* can be interpreted as inclusive fitness.

## Author Contributions

N.T. conceived the study, designed, implemented and analysed the models, and wrote the paper. K.K. discussed the design, results and implications of the study, and commented on the manuscript at all stages.

## Acknowledgements

The authors thank Stuart A. West and his group, and Ulrich F Müller for discussion, Daniel J. van der Post, Austen R. D. Ganley, and Anthony M. Poole for help with the manuscript, and Paulien Hogeweg for inspiration. The authors wish to acknowledge the contribution of NeSI to the results of this research. New Zealand’s national compute and analytics services and team are supported by the New Zealand eScience Infrastructure (NeSI) and funded jointly by NeSI’s collaborator institutions and through the Ministry of Business, Innovation and Employment. URL http://www.nesi.org.nz

## Supplementary material

The supplementary material file containing Supplementary Texts and Fig. S1 to S8 is appended to the main manuscript file.

## 1 Supporting Texts

### 1.1 An alternative agent-based model in which coexistence between P and Q is selectively neutral

In this section, we describe an alternative agent-based model in which coexistence between P and Q is neutral with respect to cellular-level selection. In the agent-based model described in the main text, coexistence between P and Q is favoured by cellular-level selection. This is due to a specific rule about complex formation, which implies that replicators multiply fastest if both P and Q provide and receive catalysis (see Methods for details). To ascertain that this specific rule about complex formation does not critically affect results, we additionally examined an alternative model in which replicators multiply fastest even if only either P or Q provides and receives catalysis. In this model, cellular-level selection does not favour coexistence between P and Q while it still tends to maximise the multiplication rate of replicators within protocells.

In the alternative model, the reaction rate constants of complex formation are defined as a function of the 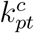 values of a replicator serving as a catalyst as follows:

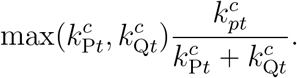

Under this definition, two replicators, denoted by *X* and *Y*, form a complex at a rate proportional to 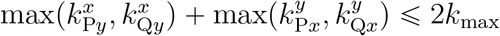 if all possible complexes are considered, where *x* and *y* are the replicator types of *X* and *Y*, respectively (in the original model, this rate is proportional to 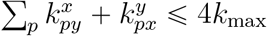. Accordingly, replicators multiply fastest not only if 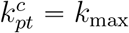 for all combinations of *c*, *p*, and *t*, but also if 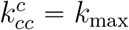 for either *c* = P or *c* Q and 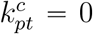 for all the other combinations of *c*, *p*, and *t*. In other words, replicators multiply fastest even if only either P or Q provides and receives catalysis (this is in contrast to the model described in the main text). While cellular-level selection always tends to maximise the multiplication rate of replicators within protocells, it is indifferent to how this maximisation is achieved. Therefore, cellular-level selection does not necessarily tend to maximise 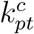 values for all combinations of *c*, *p*, and *t*; i.e., it does not necessarily favour coexistence between P and Q.

To examine the effect of coexistence between P and Q on symmetry breaking, we simulated the alternative model described above with two initial conditions, symmetric and asymmetric. In the symmetric initial condition, both P and Q were present—this is the same initial condition as used for the original agent-based model. In the asymmetric initial condition, only Q was present (see Fig. S2 for details)—this condition might be closer to what is typically imagined in the RNA world hypothesis. For both initial conditions, the model displays the same three-fold symmetry breaking as displayed by the original model (Fig. S2), indicating that the results do not depend on whether coexistence between P and Q is favoured by cellular-level selection.

### 1.2 Alternative agent-based models in which the mutation of 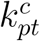 is modelled differently

In this section, we describe alternative models for the mutation of 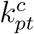. In the agent-based model described in the main text, the mutation of 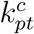 is modelled as unbiased random walks in a half-open interval (−∞, *k*_max_) with a reflecting boundary at 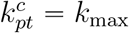. To ascertain that this specific model of mutation does not critically affect results, we additionally examined two alternative models of mutation. The first alternative model is nearly the same as the above, except that the reflecting boundary condition is set at 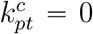. In the second alternative model, each 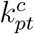 value is mutated by multiplying exp(*ϵ*), where *ϵ* is a number randomly drawn from a uniform distribution on the interval (−*δ*_mut_, *δ*_mut_), with a reflecting boundary at 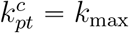. Both models of mutation produce essentially the same result as described in the main text (Figs. S3 and S4), indicating that the results do not depend on the specific models of mutation.

### 1.3 The derivation of equation (1)

In this section, we describe the derivation of equations (1) that is outlined in Methods.

To derive equations (1), we simplified the agent-based model in two ways. First, we assumed that 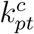 is independent of *p* and *t*. Under this assumption, a catalyst does not distinguish the replicator types of templates (i.e., 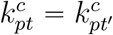 for *t* ≠ *t*′) and products (i.e., 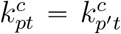 for *p* ≠ *p*′). This assumption excludes the possibility of numerical symmetry breaking, but still allows catalytic and informatic symmetry breaking as described in the main text (see Results).

Second, we abstracted away chemical reactions by defining 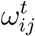 as the probability that replicator *j* of type *t* in protocell *i* is replicated or transcribed per unit time. Let 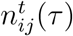 be the population size of this replicator at time *τ*. Then, the dynamics of 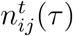 can be mathematically described as

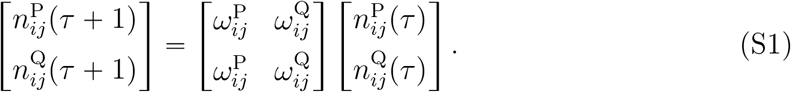

The fitness of the replicator can be defined as the dominant eigenvalue *λ*_*ij*_ of the 2 × 2 matrix on the right-hand side of equation (S1). The equilibrium frequencies of P and Q are given by the right eigenvector ***v***_*ij*_ associated with *λ_ij_*. Fisher’s reproductive values of P and Q are given by the corresponding left eigenvector ***u***_*ij*_. These eigenvalue and eigenvectors are calculated as follows:

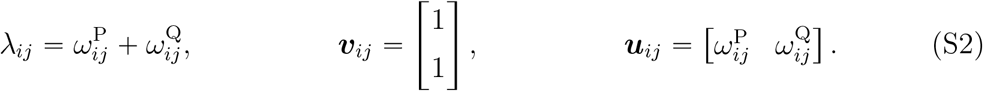

Based on the above simplification, we now derive equations (1). For concreteness, we focus on the evolution of the average catalytic activity of P (denoted by 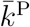 in the main text). However, the same method of derivation is applicable to that of Q if P and Q are swapped.

Let 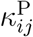 be the catalytic activity of replicator *j* of type P in protocell *i* (we use *κ* instead of *k* to distinguish 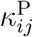 from 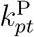). Price’s equation [1, 2] states that

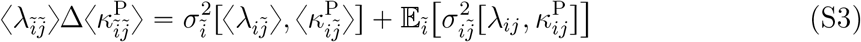

where 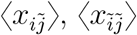, and 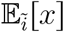 are *x* averaged over the indices marked with tildes, 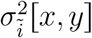 is the covariance between *x* and *y* over protocells, and 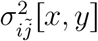 is the covariance between *x* and *y* over the replicators in protocell *i* (one replicator is always counted as one sample in calculating all moments). Below, we show that equation (S3) is approximated by equations (1) up to the second moments of 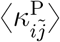 and 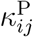, namely, 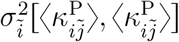 and 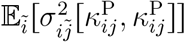.

To approximate the first term on the right-hand side of equation (S3), we assume that 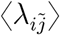 is a function of 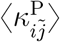 and 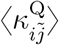 that can be expanded as a Taylor series around 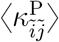 and 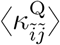. Substituting this series into 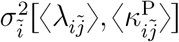, we obtain

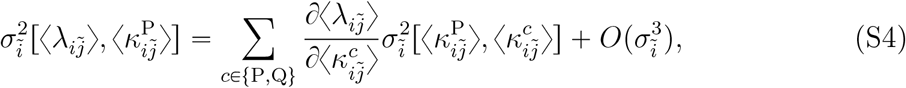

where 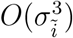 consists of terms involving the third or higher (mixed) central moments of 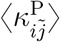 and 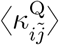 over protocells [3].

To approximate the second term on the right-hand side of equation (S3), we likewise assume that *λ_ij_* is a function of 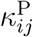 and 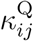 that can be expanded as a Taylor series around 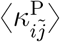 and 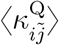. Substituting this series into 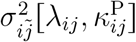, we obtain

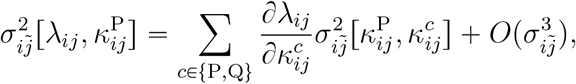

where 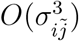 consists of terms involving the third or higher (mixed) central moments of 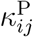 and 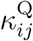 over the replicators in protocell *i* [3]. Applying 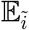 to both sides of the above equation and assuming that 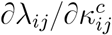 is independent of 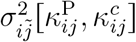, we obtain

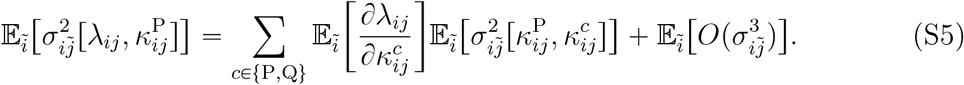

Substituting equations (S4) and (S5) into equation (S3), we obtain

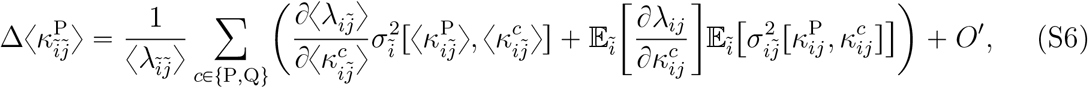

where 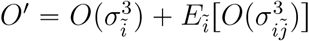.

Next, we assume that covariances 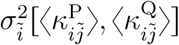 and 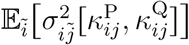 are negligible because the mutation of 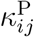 and that of 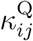 uncorrelated in the simulation model (this assumption is alternatively justified in SI Text 1.6). Under this assumption, equation (S6) is transformed into

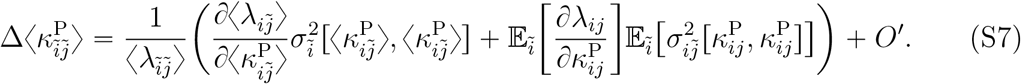

Using equation (S2) (i.e., 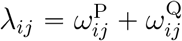), we can transform equation (S7) into

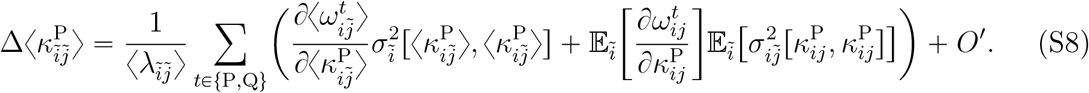

Moreover, it can be shown that

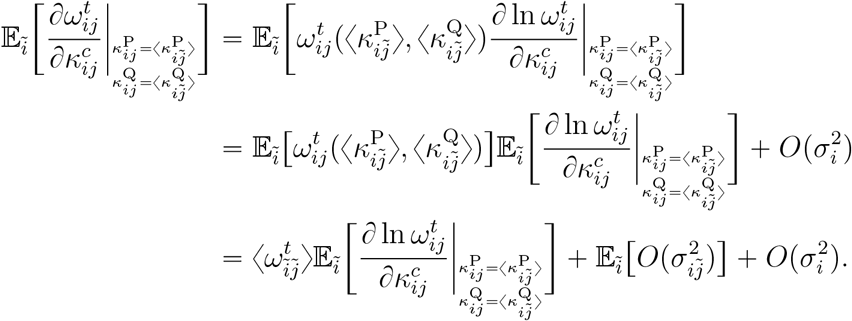

Using the above equation, we can transform equation (S8) into

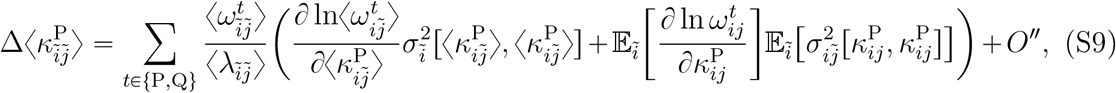

where 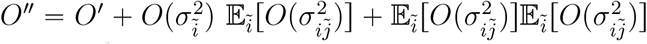.

We adopt the following notation:

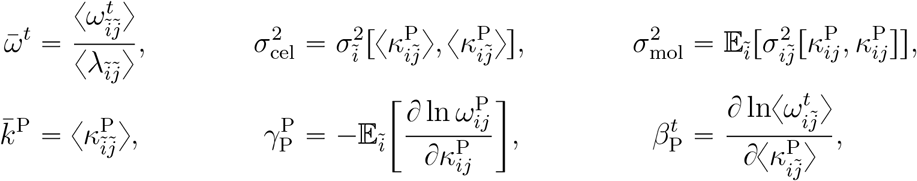

where 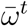 is the normalised average reproductive value of type-*t* replicators, 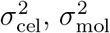, and 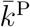 are the simplification of the notation, 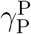 is an average decrease in the replication rate of a type-P replicator due to an increase in its own catalytic activity, and 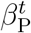 is an increase in the average replication rate of type-*t* replicators in a protocell due to an increase in the average catalytic activity of type-P replicators in that protocell.

We assume that *V* is so large that 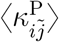 and 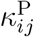 can be regarded as mathematically independent of each other, provided *i* and *j* are fixed (if *i* and *j* are varied, 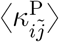 and 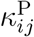 may be statistically correlated). Under this assumption, increasing 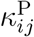 does not increase 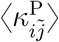, so that 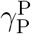 reflects only the cost of providing catalysis at the molecular level. Likewise, increasing 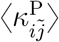 does not increase 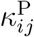, so that 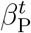 reflects only the benefit of receiving catalysis at the cellular level. Moreover, the independence of 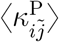 from 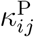 implies that 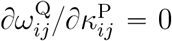, which permits the following interpretation: if a replicator of type P provides more catalysis, its transcripts, which is of type Q, pay no extra cost (i.e., 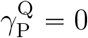).

Using the above notation and the fact that 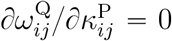, we can transform equation (S9) into

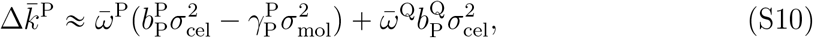

where *O*″ is omitted. equation (S10) is identical to equations (1).

Finally, to derive the equation for 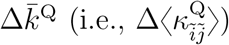, we swap P and Q in the above derivation. Moreover, we assume that 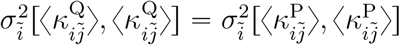 and 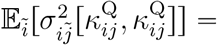 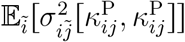 because no difference is a priori assumed between P and Q.

### 1.4 The mathematical analysis of numerical symmetry breaking

In this section, we show that numerical symmetry breaking occurs because while it is neither favoured nor disfavoured by molecular-level selection, it is favoured by cellular-level selection if catalytic and informatic symmetry breaking has occurred. To this end, we will again simplify the agent-based model into mathematical equations in a mannar analogous to that used to derive equations (1).

Before describing the mathematical analysis, we first need to note that the proximate—as opposed to ultimate—cause of numerical symmetry breaking is the self-replication of catalysts (i.e., 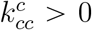, where *c* is the replicator type of catalysts) in the absence of the reverse transcription of catalysts (i.e., 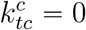, where *t* is the replicator type of templates). This fact can be inferred from the following two results. First, when catalytic, informatic, and numerical symmetry breaking occurs, the replication and transcription of templates are catalysed at about the same rate, i.e., 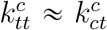 (Fig. 2b). Therefore, the replication and transcription of templates cannot cause numerical asymmetry. Second, when catalytic and informatic symmetry breaking occurs without numerical symmetry breaking, the self-replication of catalysts is absent (Fig. S5). Taken together, these results indicate that the proximate cause of numerical symmetry breaking is the self-replication of catalysts in the absence of the reverse transcription of catalysts. Therefore, to understand why numerical symmetry breaking occurs, we need to understand why the self-replication of catalysts evolves if catalytic and informatic symmetry breaking has occurred.

To address the above question, we assume that replicators have already undergone catalytic and informatic symmetry breaking and consider how the fitness of those replicators depends on the self-replication of catalysts. The population dynamics of replicators with catalytic and informatic asymmetry can be described as follows. Let 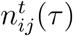 be the population size of replicator *j* of type *t* in protocell *i* at time *τ*. Let catalysts and templates be P and Q, respectively. Then, the dynamics of 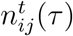 is mathematically described as follows:

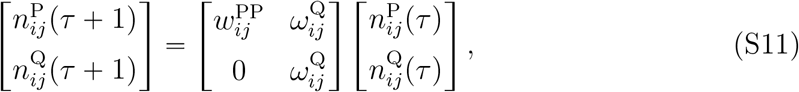

where 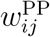 is the self-replication probability of catalysts, and 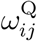 is the replication and transcription probabilities of templates, which are assumed to be identical to each other. The fitness of replicators can be defined as the dominant eigenvalue (denoted by *λ_ij_*) of the 2 × 2 matrix on the right-hand side of equation (S11):

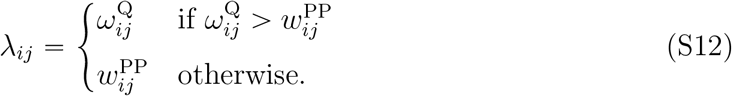

The associated right eigenvector, which determines the stationary frequencies of P and Q, is

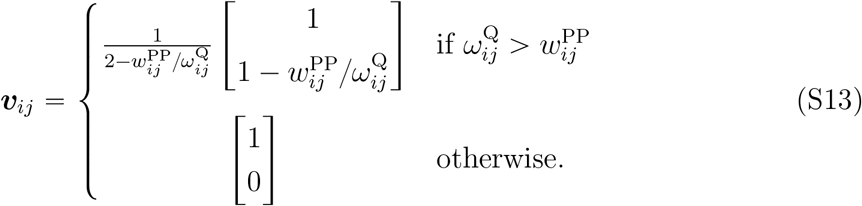

Equation (S13) shows that we must assume 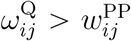 in order for P and Q to coexist. Equation (S13) also shows that the frequency of catalysts (i.e., 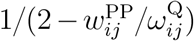) increases with the self-replication of catalysts (i.e., 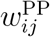), as stated in the beginning of this section.

We first examine whether the self-replication of catalysts is favoured by molecular-level selection. To this end, we consider how the fitness of replicators (i.e., *λ_ij_*) depends on the self-replication of catalysts (i.e., 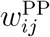). According to equation (S12), *λ_ij_* does not directly depend on 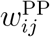. However, *λ_ij_* can indirectly depend on 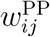 because *λ_ij_* increases with the frequency of catalysts in a protocell (i.e., 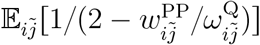). This frequency increases with 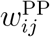 if *V* is so small that a particular replicator can influence the frequency of catalysts in the protocell. However, if *λ_ij_* increases with 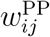, the average fitness of replicators in the protocell (i.e., 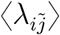) must also increase. Therefore, we need to consider the relative fitness (i.e., 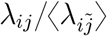). The relative fitness is independent of 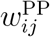 because catalysis is equally shared among templates within a protocell. Therefore, the self-replication of catalysts is neither favoured not disfavoured by molecular-level selection.

We next examine whether the self-replication of catalysts is favoured by cellular-level selection. To this end, we consider how the fitness of a protocell depends on the average self-replication of catalysts in that protocell (i.e., 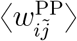). The fitness of a protocell can be defined as the average fitness of the replicators in that protocell (i.e., 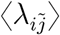). According to equation (S12), 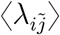 does not directly depend 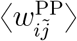. However, 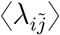 indirectly depends on 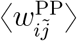 because 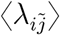 increases with the frequency of catalysts in a protocell (i.e., 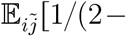 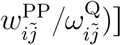). This frequency increases with 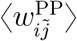, so that 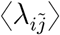 must also increase with 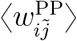. Therefore, the self-replication of catalysts is favoured by cellular-level selection.

Taken together, the above considerations indicate that the self-replication of catalysts is neutral with respect to molecular-level selection, but advantageous with respect to cellular-level selection. Therefore, numerical symmetry breaking results from the maximisation of fitness at the cellular level in the presence of catalytic and informatic asymmetry.

Finally, we mention an important consequence of numerical symmetry breaking. Numerical symmetry breaking causes a bottleneck effect on the population of replicators within a protocell. This bottleneck effect increases among-cell variance relative to within-cell variance (i.e., 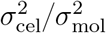); therefore, it has a stabilising effect on protocells [4, 5]. In this regard, numerical symmetry breaking can be compared to life-cycle bottlenecks displayed by multicellular organisms and eusocial colonies (i.e., an organism or colony develops from only one or a few propagules), which are considered to reduce within-group conflict [6–8].

### 1.5 The hierarchical Wright-Fisher model

In this section, we describe a model that stochastically simulates the population dynamics described by equations (1), in which 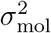 and 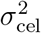 are treated as dynamic vdaeprieanbdleesnt on *m* and *V*.

The simplifications involved in the derivation of equations (1), while illuminating, make the comparison between equations (1) and the agent-based model indirect. Specifically, equations (1) cannot be compared with the agent-based model in terms of the same parameters, because the equations treat 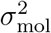 and 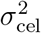 as parameters, which are adcytnuaamllyic variables dependent on *m* and *V* in the agent-based model. To fill this gap, we constructed a model that stochastically simulates the population dynamics described by equations (1) and treats 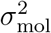 and 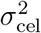 as dynamic variables dependent on *m* and *V*.

This model is formulated as a hierarchical Wright-Fisher process. Replicators are partitioned into a number of groups (hereafter, protocells). Each replicator is individually assigned replicator type *c* ∈ {P, Q} and two *k^c^* values. The fitness of a replicator is calculated according to equation (S14). In each generation, replicators are replicated or transcribed with probabilities proportional to 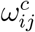, so that the population dynamics matches equation (S1) on average. After the replication-transcription step, the proto-cells containing greater than *V* replicators are divided with their replicators randomly distributed between the two daughter cells. The protocells containing no replicators are discarded.

The mutation of *k^c^* is modelled as unbiased random walks with reflecting boundaries. That is, each *k^c^* value of a replicator is mutated with a probability *m* per replication or transcription by adding a number randomly drawn from a uniform distribution on the interval (−*δ*_mut_, *δ*_mut_) ((*δ*_mut_ = 0.1). The values of *k^c^* are bounded in 0, 1 with reflecting boundaries at both bounds.

To determine the condition for symmetry breaking, we simulated the above Wright-Fisher model for various values of *V* and *m*. The simulations show that symmetry breaking occurs only if *V* and *m* are sufficiently large (Fig. S8), a result that is consistent with the outcomes of the original agent-based model (Fig. 2). Given that the Wright-Fisher model involves many of the simplifications involved in equations (1), the above consistency supports the validity of the symmetry breaking mechanism described by equations (1).

### 1.6 The phase-plane analysis

In this section, we describe the phase-plane analysis outlined in Methods.

To perform the phase-plane analysis depicted in Fig. 3, we adapted equations (1) by defining 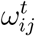 as a specific function of 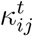 (see the previous section for the meaning of 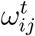 and 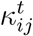). The following definition was employed:

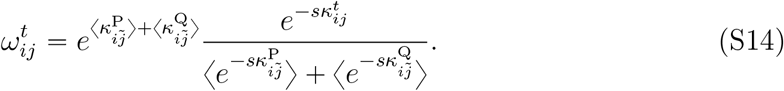

where the factor 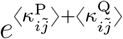 represents the cellular-level benefit of catalysis provided by the replicators in protocell *i*, the numerator 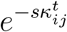 represents the molecular-level cost of catalysis provided by the focal replicator, the denominator 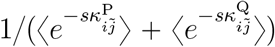 normalises the cost, and *s* is the cost-benefit ratio. The above definition of 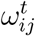 was chosen to satisfy the requirement that a replicator faces the trade-off between providing catalysis and serving as a template, so that 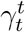 and 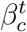 are positive; for example, if the cost 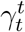 were negative, it would actually be a benefit, so that there would be no trade-off. This requirement is satisfied if 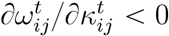 and 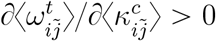 for *c* = *t* and *c* ≠ *t*. Apart from this requirement, the definition was arbitrarily chosen for simplicity.

Under the definition of 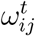 in equation (S14), we obtain equations describing the evolution of 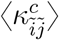 (denoted as 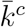 in the main text) as follows. Since the evolution of 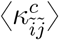 is described by equation (S6), we substitute equation (S14) into equation (S6). For this substitution, we need to calculate the derivatives of fitness. According to equation (S2), the fitness of a replicator is 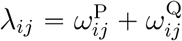. Therefore,

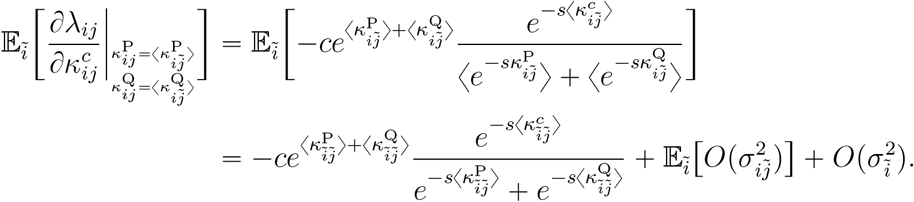

Moreover, the average fitness of replicators in a protocell is 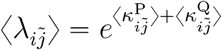, so

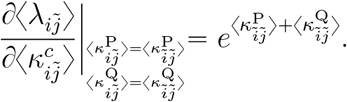

We substitute these derivatives into equation (S6) and use the fact that

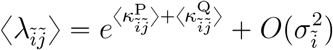

to obtain

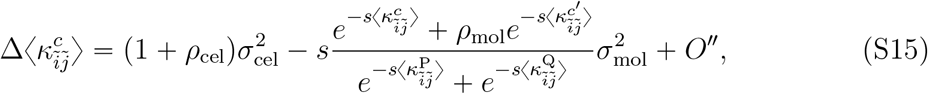

where *c*′ ≠ *c*, *ρ*_cel_ is the correlation coefficient between 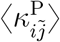 and 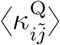 (i.e., 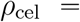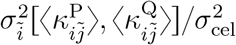), and *ρ*_mol_ is the average correlation coefficient between 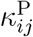 and 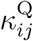 (i.e., 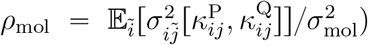). To derive equation (S15), we have assumed that the variances of 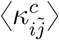 and 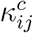 are independent of *c*; i.e., 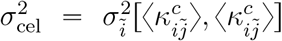 and 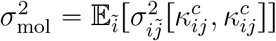 for *c* = P and *c* = Q.

Equation (S15) can be expressed in a compact form as follows:

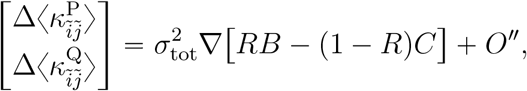

where ∇ is a nabla operator (i.e., 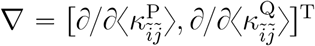, where ^T^ denotes transpose), 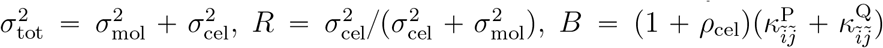, and 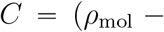 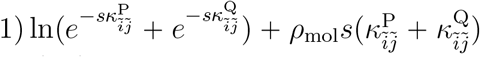. *R* can be interpreted as the regression coefficient of 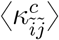 on 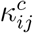 [9] and, therefore, the coefficient of genetic relatedness [10]. The potential function *RB* − (1 − *R*)*C* can then be interpreted as inclusive fitness.

Next, we set *ρ*_mol_ = 0 and *ρ*_cel_ = 0 in equations (S15) and let 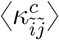 be denoted by 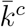, obtaining

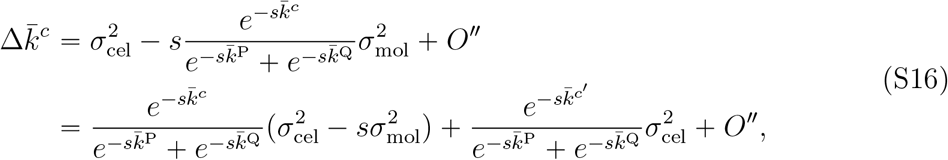

where *c*′ ≠ *c*. Comparing equations (S16) and (S10), we infer that

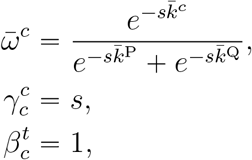

which are identical to equations (5).

Next, we omit *O*″ in equation (S16) and replace Δ with time derivative *d*{*dτ*, obtaining

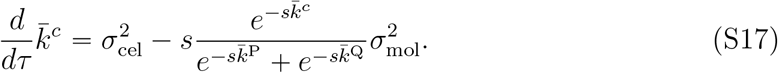

Finally, to allow for the restriction on the range of 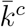 (i.e., 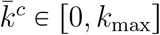), we multiply the right-hand side of equation (S17) with a function, denoted by 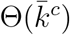, that is 1 if 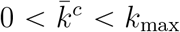 and 0 if 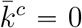 or 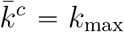. Multiplying 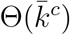 with the right-hand side of equation (S17), we obtain

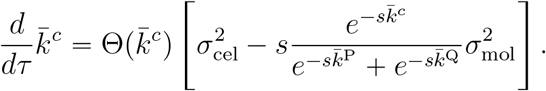

The above equation was numerically integrated for *s* = 1 to obtain the phase-plane portrait depicted in Fig. 3.

Equation (S15) allows for statistical correlations between 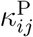 and 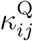 at the molecular and cellular levels, i.e., *ρ*_mol_ and *ρ*_cel_. Therefore, it can be used to examine the consequence of ignoring these correlations, which is one of the simplifications made in the derivation of equations (1) described in SI Text 1.3. For this sake, we calculate the nullcline of 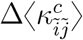. Setting 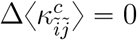 in equation (S15) and omitting *O*″, we obtain

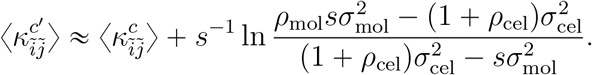

This equation shows that all parameters only appear in the intercept of the nullcline with the 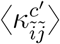-axis. Let us denote this intercept as *s*^−1^ ln *I*. The way *I* qualitatively depends on 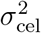 and 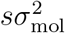 independent of *ρ*_cel_ because −1 < *ρ*_cel_ < 1. Therefore, we can assume that *ρ*_cel_ “ 0 without loss of generality. Next, to see how *ρ*_mol_ influences *I*, we focus on the singularity of *I* by setting 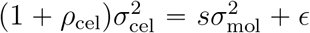, where *ϵ* > 0. Then, 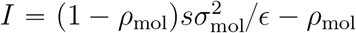. The way *I* qualitatively depends on 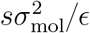 is independent of *ρ*_mol_ because −1 < *ρ*_mol_ < 1. Therefore, we can assume that *ρ*_mol_ = 0 without loss of generality. Taken together, these calculations show that ignoring correlations between 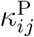 and 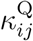 does not qualitatively affect the results, supporting the validity of equations (1).

## 2 Supporting Figures

**Figure S1:**
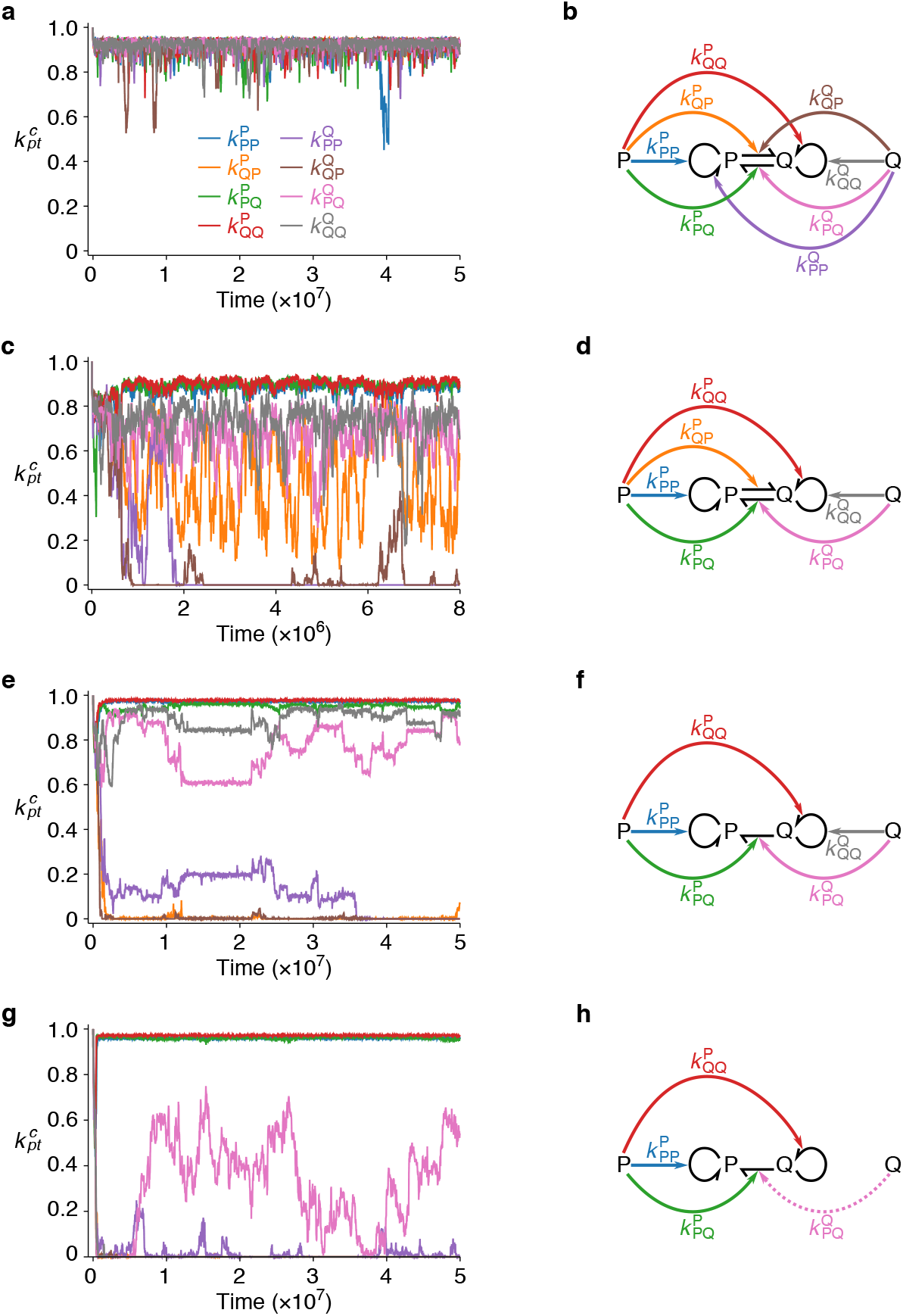
The evolutionary dynamics of the agent-based model. **a**, The dynamics of 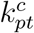 averaged over all replicators for parameters corresponding to ‘no symmetry breaking’ in Fig. 2a: *V* = 178 and *m* = 0.01. **b**, Catalytic activities evolved in a. **c**, **d**, Parameters corresponding to ‘uncategorised’ in Fig. 2a: *V* = 178 and *m* = 0.1. **e**, **f**, Parameters corresponding to ‘incomplete symmetry breaking’ in Fig. 2a: *V* = 562 and *m* = 0.01. **g**, **h**, Parameters corresponding to ‘incomplete symmetry breaking’ in Fig. 2a: *V* = 1778 and *m* = 0.01.

**Figure S2:**
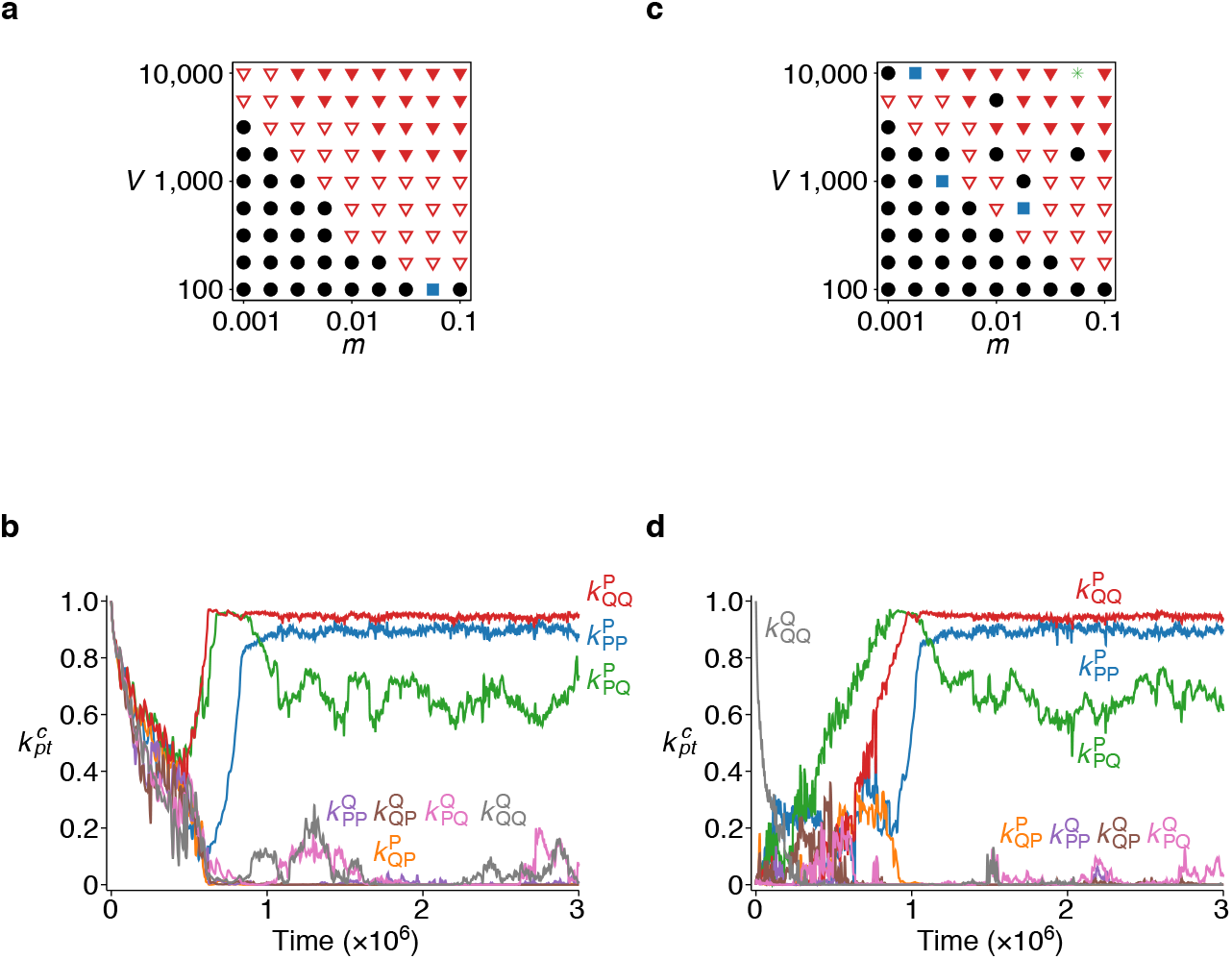
Symmetry breaking with an alternative definition of complex formation rates (see SI Text 1.1). The rate constants of complex formation were defined in such a way that coexistence between P and Q is neither favoured nor disfavoured by cellular-level selection. **a**, Phase diagram with a symmetric initial condition: 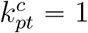 for all combinations of *c*, *p*, and *t*, with both P and Q present at the beginning of each simulation. The symbols are the same as in Fig. 2a, except that the circles include cases in which one replicator type goes extinct. **b**, Dynamics of 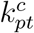 averaged over all replicators for *m* = 0.01 and *V* = 10000 in a. **c**, Phase diagram with an asymmetric initial condition: 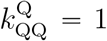 and 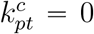 for all the other combinations of *c*, *p*, and *t*, with only Q present at the beginning of each simulation. The symbols are the same as in a, except that stars indicate the extinction of replicators. **d** Dynamics of 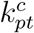 averaged over all replicators for *m* = 0.01 and *V* = 10000 in b.

**Figure S3:**
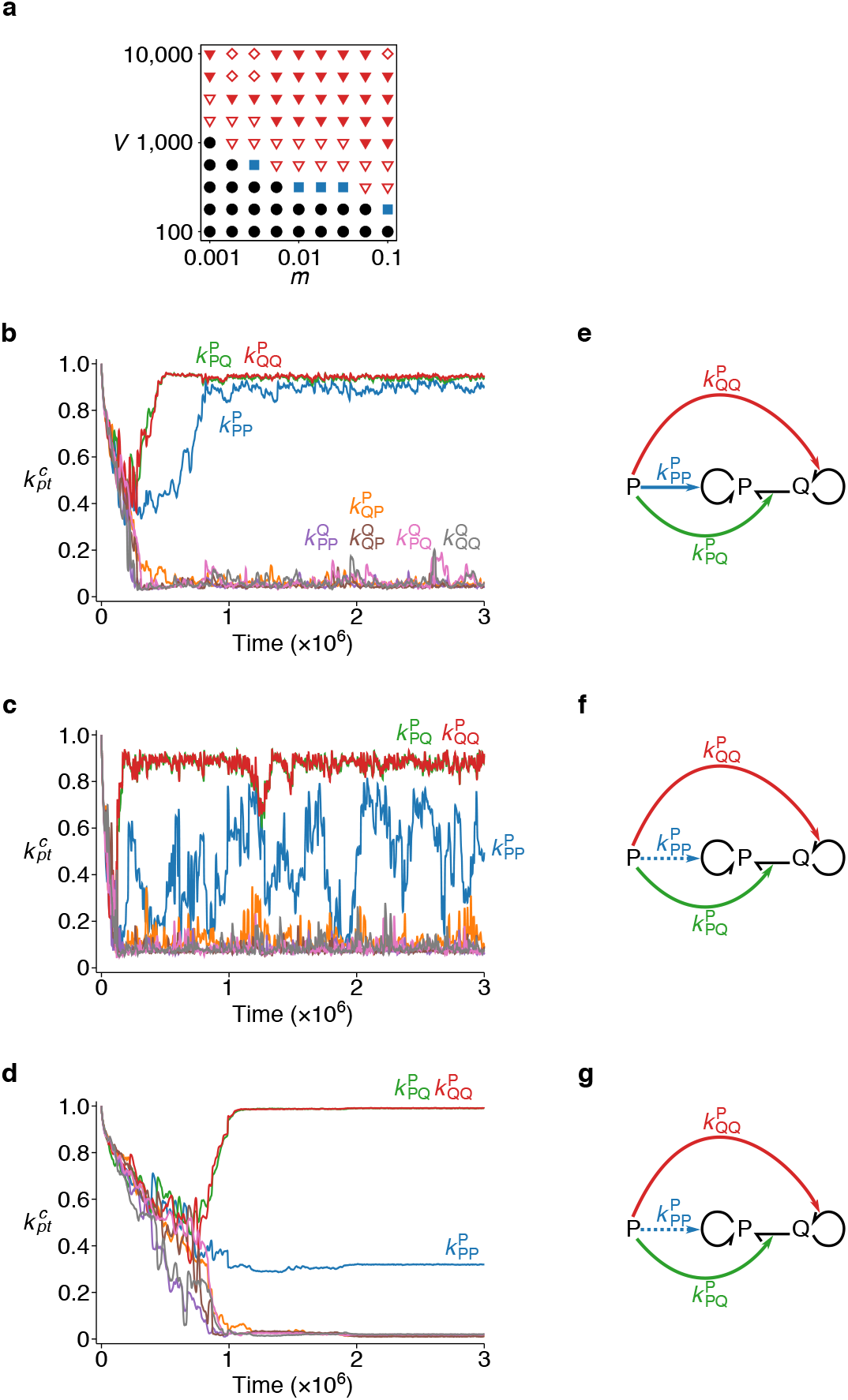
Symmetry breaking with reflecting mutation (see SI Text 1.2). The mutation of 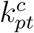 is modelled as unbiased random walk with reflecting boundaries at 0 and 1. **a**, Phase diagram. The symbols are the same as in Fig. 2a (*t*_min_ > 3.9 × 10^7^ for *m* = 0.1 and *V* = 10000). **b** Dynamics of 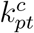 averaged over all replicators. *m* = 0.01 and *V* = 10000. Three-fold symmetry breaking occurs. **c**, *m* = 0.0562 and *V* = 10000. Numerical symmetry breaking is slight. **d**, *m* = 0.00178 and *V* = 10000. Numerical symmetry breaking is slight. **e**, **f**, **g**, Catalytic activities evolved in b, c, d, respectively.

**Figure S4:**
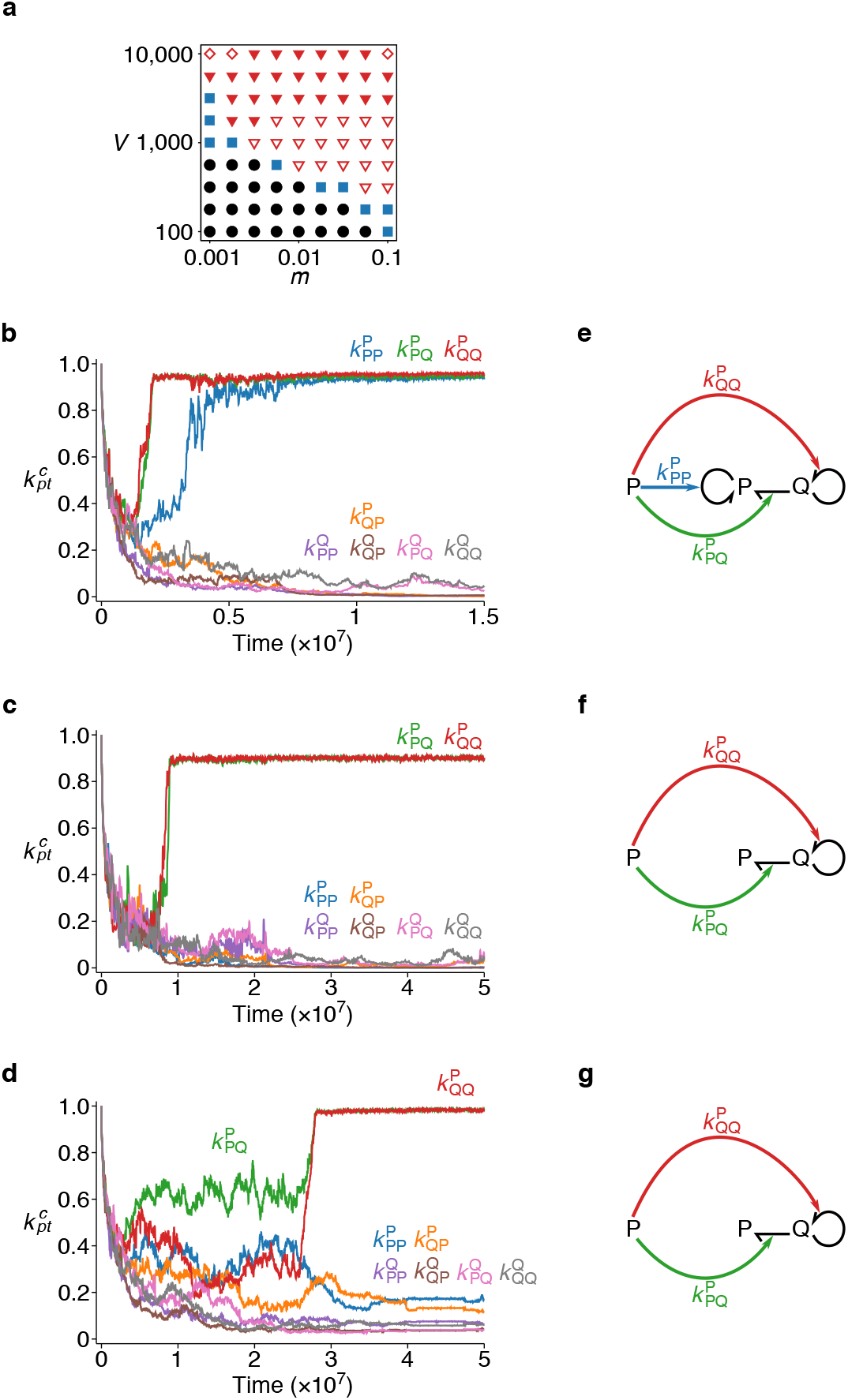
Symmetry breaking with log-space mutation (see SI Text 1.2). The mutation of 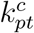 is modelled as unbiased random walks on a logarithmic scale. **a**, Phase diagram. The symbols are the same as in Fig. 2a (*t*_min_ > 3.9 × 10^7^ only for *m* = 0.1 and *V* = 10000). **b**, Dynamics of 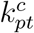 averaged over all replicators. *m* = 0.01 and *V* = 10000. Three-fold symmetry breaking occurs. **c**, *m* = 0.1 and *V* = 10000. No numerical symmetry breaking occurs. **d**, *m* = 0.00178 and *V* = 10000. No numerical symmetry breaking occurs. **e**, **f**, **g**, Catalytic activities evolved in b, c, d, respectively.

**Figure S5:**
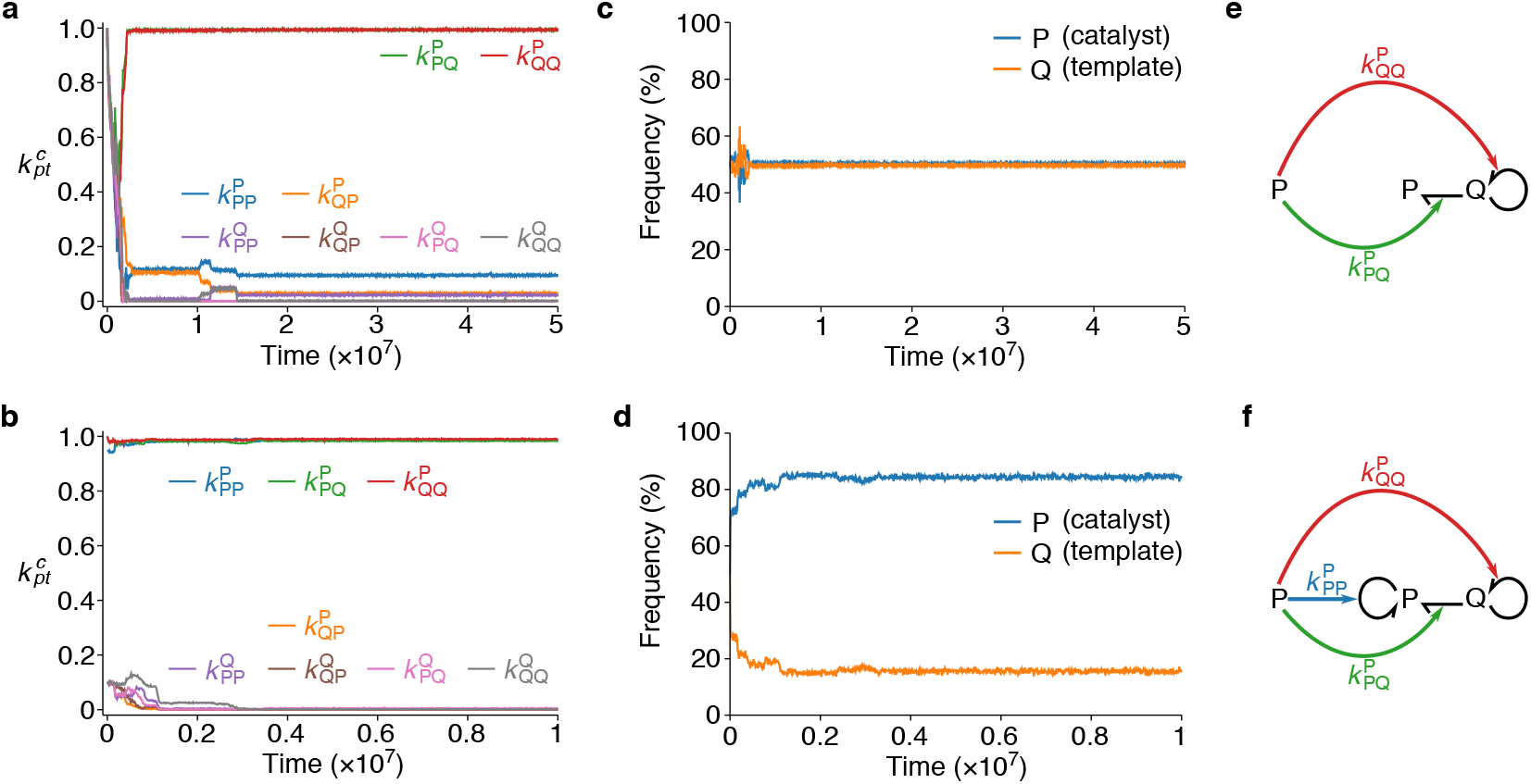
The absence of numerical symmetry breaking for small *m* and large *V* (see SI Text 1.4). **a**, **b**, The dynamics of 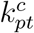 averaged over all replicators is shown for *V* = 10000 and *m* = 0.001 with two different initial conditions: a symmetric initial condition, where 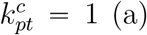 (a); an asymmetric initial condition, where 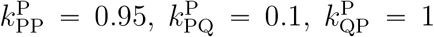, 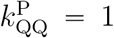, and 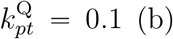 (b). The self-replication of catalysts does not evolve for the symmetric initial condition, whereas it is maintained for the asymmetric initial condition (*t*_min_ > 1.2 × 10^7^). The dependence of the results on the initial conditions suggests the presence of bistability for *V* = 10000 and *m* = 0.001. **c**, **d**, The frequencies of P (catalysts) and Q (templates) are plotted as the functions of time. Numerical symmetry breaking does not occur for the symmetric initial condition, whereas it occurs for the asymmetric initial condition. The results indicate that numerical asymmetry depends on the self-replication of catalysts. **e**, **f**, Catalytic activities evolved for the symmetric initial condition (e) and for the asymmetric initial condition (f).

**Figure S6:**
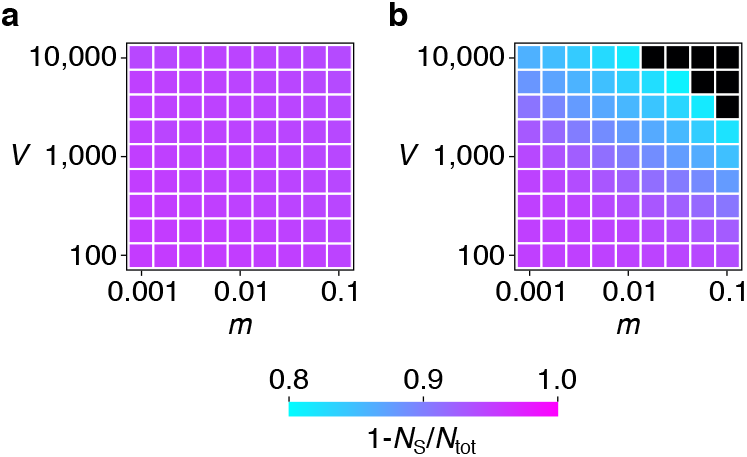
The effect of symmetry breaking on catalytic activities. The fraction of replicators 1 − *N*_S_/*N*_tot_, which is a proxy for the overall catalytic activity of replicators, is shown as a function of *m* and *V*, where *N*_S_ is the total number of S molecules in the system, and *N*_tot_ = *N*_P_ + *N*_Q_ + *N*_S_. **a**, The original model, which allows symmetry breaking (i.e., Fig. 1). **b**, The model that excludes the possibility of symmetry breaking; specifically, it allows only one type of replicator (either P or Q). Black squares indicate extinction (i.e. *N*_tot_ = *N*_S_). *t*_min_ > 1.5 × 10^7^.

**Figure S7:**
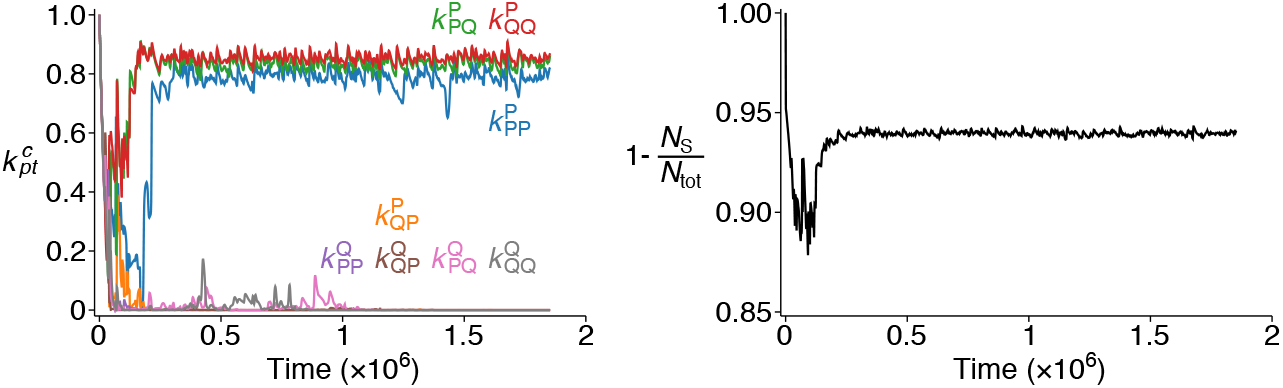
Result for large *m* and *V* values. The dynamics of the agent-based model is shown for *m* = 0.1 and *V* = 10^5^, parameters outside the range examined in Fig. 2a and Fig. S6a. **a**, The dynamics of 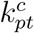 averaged over all replicators. **b**, The dynamics of the fraction of replicators 1 − *N*_S_/*N*_tot_, where *N*_tot_ and *N*_S_ are the total numbers of particles and S molecules in the system, respectively. *t*_min_ > 1.8 × 10^6^.

**Figure S8:**
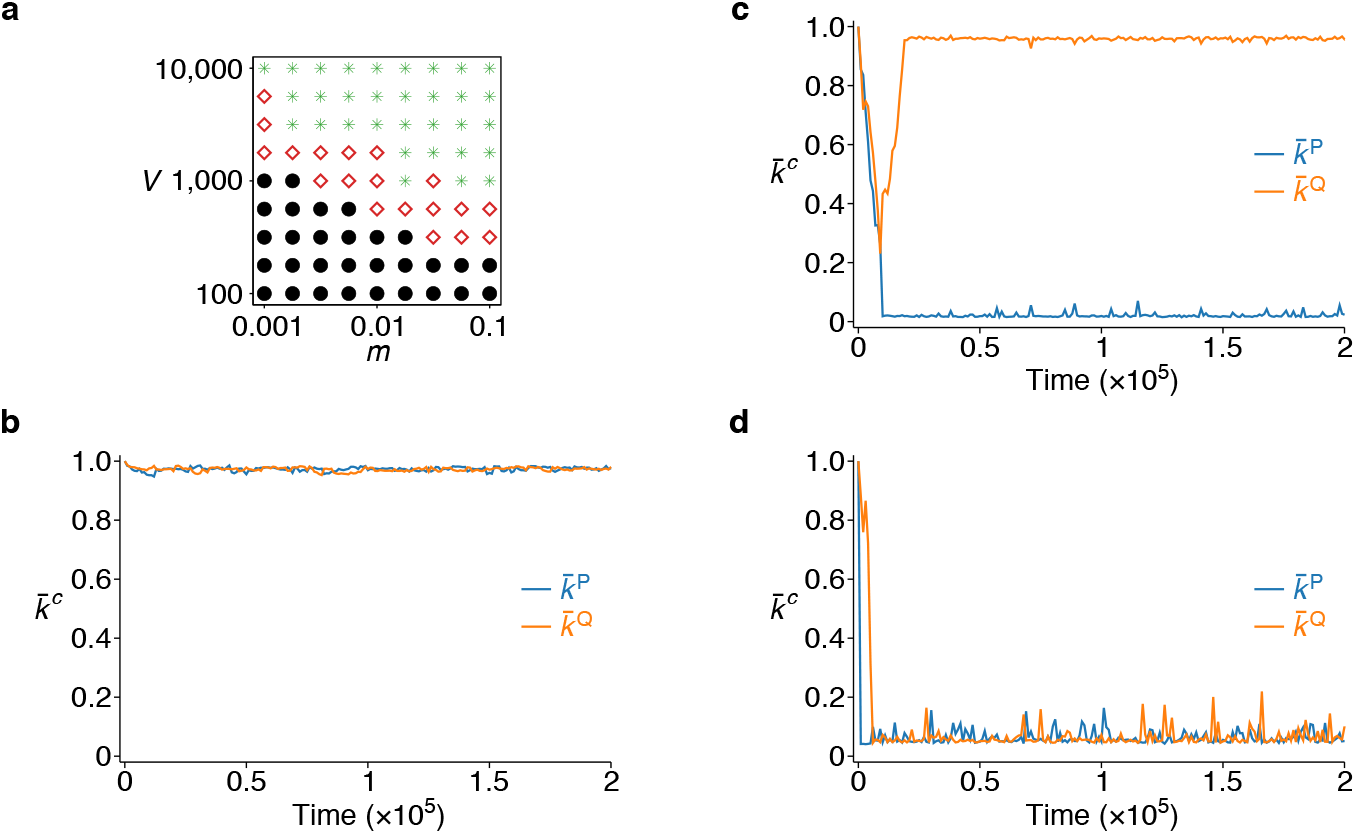
Symmetry breaking in a hierarchical Wright-Fisher model (see SI Text 1.5). The model stochastically simulates the population dynamics described by equations (1), treating 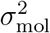 and 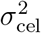 as variables dependent on *m* and *V* (see SI Text 1.5). **a**, Phase diagram. Circles indicate no symmetry breaking (i.e., 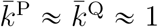); diamonds, symmetry breaking (i.e., 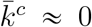 and 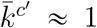 for *c* ≠ *c*′); stars, extinction (i.e., 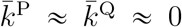). *s* = 1 (cost-benefit ratio). The total number of replicators was 50*V* (approximately 130 protocells throughout simulations). The initial condition was *k*^P^ = *k*^Q^ = 1 for all replicators. Each simulation was run for 4 × 10^5^ generations. The extinction (i.e., 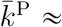 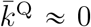) for large *m* and *V* is consistent with the phase-plane analysis of equations (1), which also shows extinction (i.e., 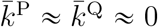) for sufficiently large 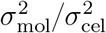 (parameters outside the range examined in Fig. 3). The discrepancy between Fig. S8a and Fig. 2a is due the simplifying assumption made in equations (1) that 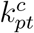 is independent of *p* and *t*. If 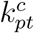 is allowed to depend on *p* and *t*, the flow of information from templates to catalysts can become completely unidirectional. Such unidirectional flow of information can resolve the dilemma between catalysing and templating and leads to the maintenance of high catalytic activities as described in Results. **b**, The dynamics of 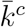 for *m* = 0.001 and *V* = 1000 (no symmetry breaking). **c**, *m* = 0.01 and *V* = 1000 (symmetry breaking). **d**, *m* = 0.1 and *V* = 1000 (extinction).

